# CorRelator: An interactive and flexible toolkit for high-precision cryo-correlative light and electron microscopy

**DOI:** 10.1101/2020.08.06.240481

**Authors:** Jie E. Yang, Matthew R. Larson, Bryan S. Sibert, Samantha Shrum, Elizabeth R. Wright

## Abstract

Cryo-correlative light and electron microscopy (CLEM) is a technique that uses the spatiotemporal cues from fluorescence light microscopy (FLM) to investigate the high-resolution ultrastructure of biological samples by cryo-electron microscopy (cryo-EM). Cryo-CLEM provides advantages for identifying and distinguishing fluorescently labeled proteins, macromolecular complexes, and organelles from the cellular environment. Challenges remain on how correlation workflows and software tools are implemented on different microscope platforms to support microscopy-driven structural studies. Here, we present an open-source desktop application tool, CorRelator, to bridge between cryo-FLM and cryo-EM/ET data collection instruments. CorRelator was designed to be flexible for both on-the-fly and post-acquisition correlation schemes. The CorRelator workflow is easily adapted to any fluorescence and transmission electron microscope (TEM) system configuration. CorRelator was benchmarked under cryogenic and ambient temperature conditions using several FLM and TEM instruments, demonstrating that CorRelator is a rapid and efficient application for image and position registration in CLEM studies. CorRelator is a cross-platform software featuring an intuitive Graphical User Interface (GUI) that guides the user through the correlation process. CorRelator source code is available at: https://github.com/wright-cemrc-projects/corr.

## Introduction

Cryo-correlative light and electron microscopy (cryo-CLEM) is the coupling of (cryo-) fluorescence light microscopy (FLM) and cryo-electron microscopy (cryo-EM). Cryo-CLEM has proven to be a powerful method for *in situ* structural studies, including time-dependent events, especially those associated with viral entry, replication, and egress (Briegel et al., 2010; Fu et al., 2019; Hampton et al., 2016; Jun et al., 2019; Koning et al., 2014; Koster & Grünewald, 2014; Schorb et al., 2017; Wolff et al., 2016; Zhang, 2013). Fluorescent labeling and light-level imaging of viral and cellular factors enables one to capture dynamic events during virus-infection of cultured cells for real time rapid identification of targets, and for orienting on their spatial positions. Subsequently, cryo-electron microscopy (cryo-EM) and tomography (cryo-ET) are used to determine nanometer- and sub-nanometer-resolution three-dimensional (3D) structures of macromolecules present in virus-infected cells (Bharat et al., 2017; Brandt et al., 2010; Briggs, 2013; Dick et al., 2020; Erlendsson et al., 2020; Kovtun et al., 2018; Strauss et al., 2015; Wan et al., 2017). In cryo-EM, the rapid vitrification of biological samples minimizes preparative artifacts typically associated with conventional EM procedures such as chemical fixation, dehydration, and resin-embedding (Hampton et al., 2016; Faas et al., 2013; Jun et al., 2011; Lučić et al., 2013, 2007; Moser et al., 2019). This is important for *in vivo* investigations of pleomorphic viruses and cell membrane-connected events such as viral assembly and budding, because membrane morphology and membrane-associated protein organization may be altered by conventional fixation protocols (Ke, Dillard, et al., 2018; Ke, Strauss, et al., 2018; Luque & Castón, 2020).

In cryo-CLEM, sample vitrification can be performed either before or after fluorescence imaging (Briegel et al., 2010). Fluorescence imaging of a specimen prior to sample vitrification is important because of the improved resolution of immersion-objective microscopes and options for dynamic, time-resolved imaging (Fu et al., 2019; Jun et al., 2011). However, specimen states observed prior to cryo-fixation may change, be disrupted, or damaged by the vitrification process, thereby limiting the precision of correlation. Cryo-fluorescence microscopes (cryo-FLM) equipped with dedicated ‘cryo-stages’ that maintain specimens well below −150°C have made it possible to correlate biological events on vitrified samples. In recent years, various solutions have been introduced to optimize low-temperature imaging, to reduce the number of grid transfer steps, and to realize high-resolution cryo-FLM. These include the development and use of stable cryo-stages (Jun et al., 2011; Schellenberger et al., 2014; Schorb et al., 2017; Schorb & Briggs, 2014; Schwartz et al., 2007; van Driel et al., 2009), integrated cryo-FLM and −EM systems (Agronskaia et al., 2008; Faas et al., 2013; Gorelick et al., 2019; Li et al., 2018), and high numerical-aperture (NA) cryo-objective lenses (Li et al., 2018; Nahmani et al., 2017; Schorb et al., 2017). In addition, super-resolution cryo-CLEM systems have been explored, aiming to further bridge the resolution gap between LM and EM imaging modalities (Chang et al., 2014; Liu et al., 2015; Kaufmann et al., 2014; Moser et al., 2019; Nahmani et al., 2017).

Cryo-correlation typically proceeds in two steps, on-the-fly targeting and post-acquisition superposition. On-the-fly correlation guides cryo-EM/ET data collection to targets of interest (TOI), while post-acquisition transformations support precise mapping of fluorescent signals to reveal ultrastructural details of targeted or potentially unknown objects in the TEM or acquired TEM images. In order to facilitate targeting, TEM grids may be marked with numbers and letters (i.e., Finder grids) to aid in the rough correlation process. Application of fiducial markers to the sample before vitrification, such as fluorescent electron-dense microspheres, can support finer-scale correlations of approximately 10 to 100 nm (Fu et al., 2019; Kukulski et al., 2011; Li et al., 2018; Schellenberger et al., 2014; Schorb & Briggs, 2014). A less common method is to use EM grid support features, including holes and imperfections in the carbon film, to achieve marker-free correlative alignment (Anderson et al., 2018). In all cases, there are still difficulties to overcome. First, the distribution of fiducial markers across the entire grid needs to be relatively uniform for alignments to be reliable. The optimized density must be high enough to provide good alignment, but low enough not to obscure fluorescent signals from TOIs. Second, bent, warped, or crinkled vitrified grids and uneven ice-thickness hinder automated detection of the holes in the carbon films, leading to poor relocation of the same hole on both maps. Therefore, an interactive yet unbiased approach for reliable registration is needed.

A number of research groups have developed software or scripts to facilitate the coordinate transfer step (Anderson et al., 2018; Fu et al., 2019; Jun et al., 2011; Kukulski et al., 2011; Li et al., 2018; Paul-Gilloteaux et al., 2017; Rigort et al., 2010; van Driel et al., 2009; Schwartz et al., 2007). Most of the tools are available to the community, however, several are not easily incorporated into existing or under-development correlative cryo-ET data collection routines. For example, some are system-dependent and built for specific workflows and hardware configurations (Gorelick et al., 2019; Li et al., 2018). Others may require access to licensed materials such as MATLAB or for users to be savvy at programing languages (Fu et al., 2019; Kukulski et al., 2011; van Driel et al., 2009). ImageJ- or Icy-based solutions implement transformation algorithms to manage complicated geometric distortions, but are used for post-acquisition high-accuracy correlation, and are less practical for on-the-fly targeting applications. There is a growing need for specimen-specific, multi-level correlative schemes that are flexible and easily customized for current and future instruments and workflows (Sartori-Rupp et al., 2019; Strnad et al., 2015).

Here, we present a cross-platform, user-friendly desktop application tool called “CorRelator” for interactive and precise coordinate translation between cryo-FLM and cryo-EM/ET. The flexibility and system-independent nature of the tool for cryo-CLEM applications is demonstrated in combination with the microscope-control platform SerialEM (Mastronarde, 2005) and vitrified cells infected with respiratory syncytial virus (RSV). We demonstrate that efficiency and target prediction of the CorRelator cryo-CLEM workflow guide the user to select regular grid support features (e.g., hole centroids) for accurate alignment. We show that lower lateral spatial resolutions that may be present in cryo-FLM data can be partially compensated for through advanced computational background cleaning and signal deconvolution processing. To balance between unbiased automation and human intervention, CorRelator supports iterative labeling of registration positions and quick feedback-based alignment of FLM/TEM images. As more complex correlative strategies evolve and the demand for system-independent cryo-CLEM schemes increases, we believe that this tool will facilitate efforts in a broad-range of project-specific correlative workflows. CorRelator is open-source and freely available for download at (https://github.com/wright-cemrc-projects/corr).

## Materials and Methods

### Cell culture and infection on TEM grids

Quantifoil grids (200 mesh Au R2/2 and Au Finder; Quantifoil Micro Tools GmbH, Großlöbichau, Germany) were coated with an extra layer of carbon for stabilization and then glow discharged for improved hydrophilicity. The grids were subsequently coated with 100 nm and 500 nm fluorescent microspheres (TetraSpeck Microspheres, Invitrogen T7279, T7281, USA) at a 500x or 1000x dilution for 5 min, followed with a washing step using 1x PBS. HeLa cells (ATCC CCL-2, ATCC, Manassas, VA, USA) were seeded on the bead-coated grids at a density of 0.5-0.75 x 10^5^ cells/mL in glass-bottomed culture dishes (MatTek Corp., MA, USA) (Schellenberger et al., 2014). The grids and dishes seeded with HeLa cells were cultured overnight at 37 °C with 5% CO2 in DMEM complete medium supplemented with 10% fetal bovine serum (FBS), 1 μg/mL penicillin, streptomycin and amphotericin B (PSA) antibiotics. After overnight incubation, the cell-seeded grids were inoculated with the recombinant virus strain RSV rA2-mK^+^ (Hotard et al., 2012). Twenty-four hours post-infection, native immunogold labeling of RSV glycoprotein F was carried out as published previously (Yi et al., 2015), with a primary antibody (motavizumab, 4 μg/mL (gift from Larry J. Anderson, Emory University)) and secondary antibody Alexa Fluor 488 Nanogold Goat anti-human IgG (Nanoprobes, NY, USA) (Cheutin et al., 2007). 4 μl of 10 nm gold fiducial beads (Aurion Gold Nanoparticles, Electron Microscopy Sciences, PA, USA) were applied to the RSV-infected cell EM grids to aid in image alignment during the reconstruction process. The grids with RSV-infected HeLa cells were plunge-frozen using a Gatan CryoPlunge3 system with GentleBlot blotters (Gatan, Inc., Pleasanton, CA, USA).

### Correlative fluorescence microscopy

Two CLEM-CorRelator workflows were tested, using a Leica DMi8 widefield fluorescence microscope at room-temperature or a Leica EM Cryo-CLEM microscope system under cryo-conditions (Leica MicroSystems, Germany). For room-temperature correlation, TetraSpeck-coated (100 nm) Quantifoil Finder EM grids were imaged at 40 x magnification (40 x, 0.6 NA lens, dry) and 63 x magnification (63 x, 1.4 NA lens, oil-immersion) in brightfield, GFP (emission, 525 nm), and Texas Red (emission, 619 nm) channels with Micro-Manager (Edelstein et al., 2014, 2010). Cryo-FLM of the grid of RSV-infected HeLa cells was performed using a Leica EM Cryo CLEM system (50x, ceramic-tipped, 0.90 NA), through bright field and the band pass filter cubes of 525 nm and 630 nm, with the dedicated Leica LAS X microscope software (Hampton et al., 2016; Schorb et al., 2017). Images combined to generate 12 to 15 μm Z-stack projections were collected of vitrified grids at a Nyquist sampling step of 350 nm to compensate for cell thickness and wavy or warped grids. The Small Volume Computational Clearance (SVCC) implemented in the Leica LAS X THUNDER package was applied to the post-acquisition image stacks to reduce image blurring and to restore weaker or lower signals. Images and mosaic tiles were exported and used as compressed lossless RGB TIFF format. We determined the point spread function (PSF) of the fluorescent signal (emission *λ* = 525 nm) of 500 nm TetraSpeck beads in the unprocessed and SVCC-processed cryo-FLM image stacks (n = 10). Additional image processing steps such as flipping, cropping, contrast adjustment were performed in ImageJ/Fiji (Schindelin et al., 2012).

### Cryo-electron microscopy, cryo-electron tomography, and tomogram reconstruction

After FLM imaging at room temperature, the bead-coated Quantifoil Finder grids were imaged with a Tecnai T12 (ThermoScientific, Hillsboro, OR, USA) operated at 120 kV and equipped with a 4k x 4k Gatan OneView camera using SerialEM (Mastronarde, 2005). After cryo-FLM imaging, bare grids with immunolabeled RSV-infected HeLa cells were imaged using a Titan Krios (ThermoScientific, Hillsboro, OR, USA) at 300 kV. Images were acquired on a post-GIF Gatan K3 camera in EFTEM mode with a 20 eV slit. Images were recorded at magnification of 81x (4,485 Å/pixel), 470x (399 Å/pixel), 2250x (76 Å/pixel), and 19500x (4.47 Å/pixel). All grid TEM maps were collected in SerialEM.

Tilt series were collected bi-directionally with 2° increments covering a tilt range of −60 ° to 60° at a magnification of 19500x (4.47 Å/pixel) and nominal defocus of −8 μm with a total dose of 80 to 100 e^-^/Å^2^. Tilt series were aligned using 10 nm fiducial beads and reconstructed with weighted back-projection algorithm using the IMOD package (Kremer et al., 1996). The cryo-tomograms reconstructed in IMOD were denoised using the low pass Gaussian filter function on 3D volumes implemented in EMAN2 (Tang et al., 2007), followed by smooth filtering with a standard kernel of 3×3 in IMOD, to enhance contrast.

### Identification of hole centers in ImageJ/Fiji

To identify the center positions of the holes in the fluorescent TIFF images, we followed similar steps as described previously (Anderson et al., 2018). Briefly, the raw images were cropped based on targets of interest (TOI). Some images were downsized by a binning factor of 2 to increase the computational processing speed. The raw images had multiple channels including brightfield and fluorescent channels (e.g., GFP and Texas Red). The cropped fluorescent channel frame (usually with high noise) was subject to iterative non-local mean filtering (Buades et al., 2005) to optimize efficiency in hole identification in the carbon film. Alternatively, brightfield frames may be used when the shift between frames of the different channels was tolerable (less than 1% of target identification exceeding 2-pixel difference). An iterative median filter of 2 to 6 iterations and optimal threshold for binarization were applied to preserve sharpness of the hole edges. Then, the binarized images were processed with the Canny edge detector function (Canny, 1986), followed by the circular Hough transform function (Illingworth & Kittler, 1987) to determine the optimal radius for hole detection and center coordinates of each detected hole. The whole procedure was carried out using the ImageJ/Fiji platform (Schindelin et al., 2012) where the Canny Edge Detector and circular Hough Transform Functions were loaded and run as plugins. The hole center coordinates (*P_x_, P_y_*) of a fluorescent TIFF image were exported in the comma-separated value (CSV) file format. For TEM image map and montages collected in SerialEM, the hole centers were identified following the same procedure described above. SerialEM also has a built-in function to label a 2D grid of hole centers for TEM maps in its Navigator module, “Add Grid of Points” function.

### Registration of stage positions in CorRelator

Image maps are next imported into CorRelator. The default coordinate system in ImageJ/Fiji is left-handed with (0,0) defining the top left corner. Pixel coordinates in this system can be directly imported from CSV formats to an image map in CorRelator. Additional positions can be manually assigned or modified. Correlator then facilitates iteratively importing and assigning pixel positions on maps and converting pixel coordinates to the stage positions. The stage positions are exportable as a Navigator file in autodoc format that can be directly read into SerialEM. Alternatively, CorRelator can solve for an affine transformation function that directly aligns the stage positions and maps.

### Affine transformation in CorRelator

We adopted the close-form solution to the least square problem for an overdetermined system to determine the optimal transformation matrix between two modalities (Horn, 1987). While at least 4 reference coordinate pairs are required to avoid the singular matrix problem, unlimited reference points can be added and incorporated into the solution. Due to the location error in the reference pair positioning, the calculated alignment will not satisfy or fit into all pairs. Instead, it is set to find the best-fitting solution that minimizes the sum of squared errors between targets and predicted outputs.

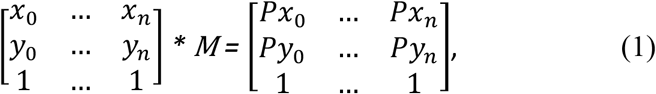

Where 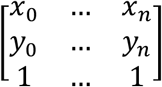 and 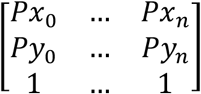 are matching homogenous reference pairs of X- and Y- image coordinates in the fluorescent image and TEM image to which the fluorescent image is registered, *M* is the transformation matrix. To simplify the equation (3), set 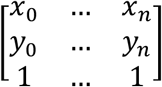 as matrix 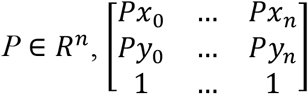 as matrix *Q* ∈ *R^n^*, then for an overdetermined system,

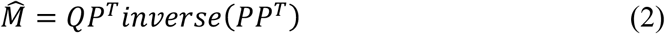

When 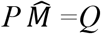 is consistent, then 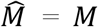 the usual matrix solution. Matrix multiplication, transposition, and inverse calculations were implemented with the Java Apache Commons Math package (Andersen et al., n.d.; Paul-Gilloteaux et al., 2017).

### Alignment accuracy estimation

We tested the alignment accuracy of CorRelator and the transformation algorithms of several programs including MATLAB (MathWorks) and eC-CLEM (Paul-Gilloteaux et al., 2017), using a procedure called “Leave-one-out” described previously (Fu et al., 2019; Kukulski et al., 2011; Schellenberger et al., 2014). Briefly, the hole centers in the carbon film within a square were used as alignment markers while either non-registered hole centroids that were unused and left out or TetraSpecks (100 or 500 nm) in the FLM image were treated as the TOIs whose positions were predicted in the registered EM image. The hole centroids were identified as described above. To select the bead signals, we used a two-dimensional Gaussian function in MATLAB. The predicted position of TOIs was then compared to the actual position to estimate the individual prediction error by measuring the Euclidean distance or length between the two points. The deviation on X and Y axes was calibrated in a form of Euclidean vector. For cryo-CLEM, Quantifoil and Quantifoil Finder grids of infected HeLa cells followed a similar procedure. In this case, F glycoprotein labeling (green signal) was treated as the TOIs. To estimate the on-the-fly image acquisition accuracy based on CorRelator’s predictions, the image shift offsets between magnifications on a SerialEM-controlled TEM were corrected to avoid additional off-target variance introduced by the microscope lens performance. We then used the deviation between TOI positions (fluorescent viruses or TetraSpecks) and the center of the acquired image after moving to the predicted stage position.

## Results

### CorRelator development

CorRelator supports both on-the-fly and post-acquisition two-dimensional (2D) cryo-correlation. The on-the-fly correlation and subsequent automated cryo-EM/ET data collection can be accomplished through SerialEM (Mastronarde, 2005), a versatile software program that controls the TEM and image detectors. While SerialEM is capable of manual point matching transformations (Hampton et al., 2016; Schorb et al., 2017; Schwartz et al., 2007), the registration process can be time-consuming and less accurate due to limited guidance and assessment steps. To tackle this challenge, CorRelator uses an iterative user-in-the-loop approach to define registration points and translate external image coordinates into adjustable TEM stage positions in SerialEM (Fig. 1 and Supplementary Fig. 1B).

**Fig. 1.**
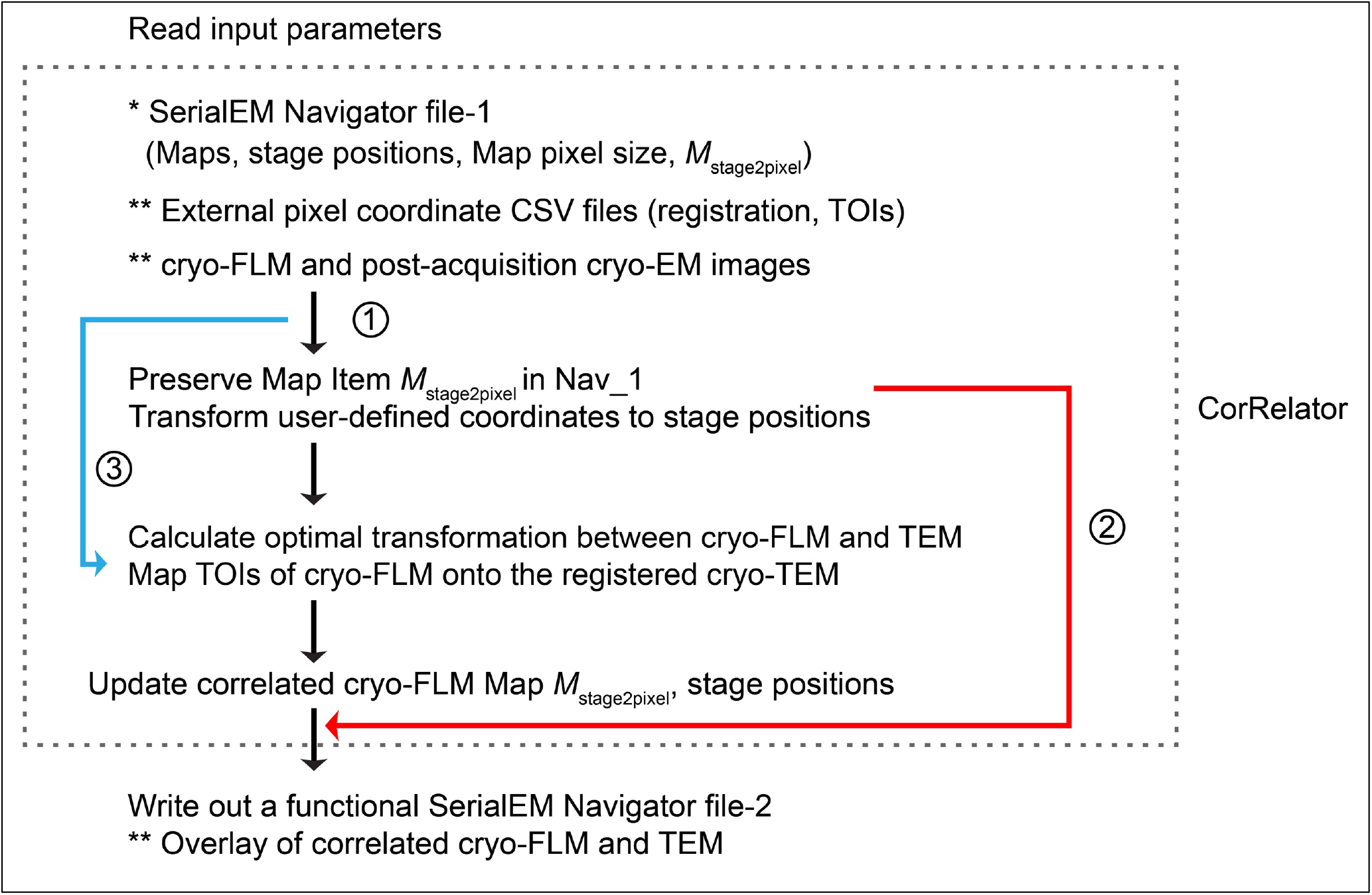
Flowchart for the algorithm implemented in the CorRelator toolkit. CorRelator supports three applications for flexible correlation, labeled as ① or ② for on-the-fly (cryo)-CLEM operation, and ③ for post-acquisition correlation. An asterisk (*) indicates the basic input file for CorRelator and ** indicative of optional inputs (e.g. independent csv files, cryo-FLM/EM frames for post-acquisition correlation) and optional outputs (e.g. overlay of correlated cryo-FLM/EM images). The dashed grey box surrounds the main operations performed in CorRelator.

The key concept of CorRelator is to use image pixel coordinates for robust registration and incorporate microscope stage-position-to-pixel-coordinate matrices to achieve fast and reliable on-the-fly correlation. We decided to use SerialEM because (1) it is an open-source program extensively used in the microscopy community; (2) it is applicable to many existing TEM systems and imaging detectors; (3) the instrumentation-determined image pixels-to-stage-position relationships are accessible in its Navigator file, where each image or mosaic is considered as a Map entry (see Appendix-A). The transformation of image planar coordinates to stage planar coordinates can be determined by a 2D affine transformation represented in homogeneous form:

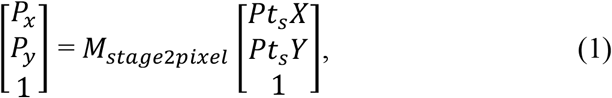

Where 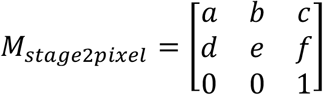 is the stage-to-pixel coordinate transform matrix, (*P_x_, P_y_*) ∈ *R*^2^ is the image planar coordinate, and (*Pt_s_X, Pt_s_Y*) ∈ *R*^2^ is the transformed TEM planar stage position. The reverse transform is:

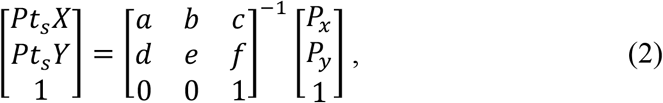

Where 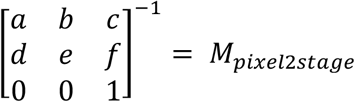 is the pixel-to-stage coordinate transform matrix.

When a set of external XY coordinates (e.g., points for registration or targets of interest, TOIs) related to a FLM or TEM map entry in the navigator file, are provided, CorRelator reads and applies the matrices *M_stage2pixel_* and *M_pixel2stage_* of the associated map to these user-defined points. As a result, the offline-determined “external” coordinates are translated to recognizable “inherent” stage positions written out in an updated Navigator file.

Vitrification ensures preservation of biological samples in a near-native state, greatly limiting physical deformations, such as non-linear warping of the sample that can occur during chemical fixation, freeze substitution, sectioning, embedding, and image acquisition. An affine transformation that preserves co-linearity of all points and ratios of distance, is suitable for accurate 2D coordinate transfer in cryo-CLEM (Fu et al., 2019; Kukulski et al., 2011; Schellenberger et al., 2014; van Driel et al., 2009). However, the registration may not be accurate for all reference pairs due to possible errors in the point pair positioning. To handle alignment in CorRelator, the transformation matrix *M* between cryo-FLM and EM is calculated using closed-form ordinary least-squares solutions in digital geometric image correlation (Castleman, 1995; Gonzalez & Woods, 2002; Horn, 1987). The transformation matrix *M* considers changes in magnification, scaling, rotation, translation, and shearing between FLM and TEM images. The transformation *M* is then applied to the coordinates of the fluorescent signal of interest, defining individual translated pixel and stage positions in the correlated TEM image.

A flow chart of the algorithm for CorRelator is shown in Figure. 1. CorRelator first defines the *M_stage2pixel_* of different maps in a baseline SerialEM navigator file (Nav_1). The map may be: (1) whole grid cryo-EM montages at low magnification with identifiable grid squares, (2) intermediate magnification mosaics of good areas, or (3) an imported image, i.e., a cryo-FLM frame. CorRelator adopts an iterative registration approach, supporting (1) interactive manual selection and or (2) the import of comma-separated values (CSV) files with pixel coordinates from external sources. A csv file is a common file format exported by many image analysis tools such as ImageJ/Fiji, IMOD, MATLAB, and other commercial software packages (Kremer et al., 1996; Schindelin et al., 2012). When provided with two matching sets of pixel coordinate pairs for image registration, CorRelator converts them into stage positions and calculates the optimal transformation matrix *M* to align the FLM to TEM frame. The alignment performance can be visually and quantitatively assessed. As a result, the user will be able to quickly decide if another round of transformation is necessary. The output is an updated functional navigator file referred as to Nav_2. Based on the experimental design and workflow, Nav_2 may be: (1) exported at various stages, (2) used for direct automated cryo-EM/ET data acquisition at the stage positions of the transformed fluorescent TOIs (Fig.1 Route 1), or (3) used for further manipulation and transformation in SerialEM (Fig. 1 Route 2). The tool is also suitable for post-acquisition correlation (Fig 1. Route 3). In this case, a matching set of x- and y-pixel coordinate pairs identifiable in FLM and EM images are direct registration points for matrix calculation.

### CorRelator graphical user interface (GUI) design and workflow

Another component of CorRelator is the intuitive graphical user interface (GUI) that assembles robust and flexible correlation paths. The software tool is a cross-platform Java/JavaFX 8 desktop application with dependencies from Apache Commons Math for matrix manipulations and Java Advanced Imaging library for graphical format support. The GUI supports multiple pathways for data management, including a *Project view* (Fig. 2 and Supplementary Fig. 1) and a *Wizard*. The *Wizard* feature (Fig. 2) is a step-by-step workflow for performing guided on-the-fly and post-acquisition cryo-FLM to EM transformation. The *Project view* alternatively allows iterative operations for importing or adding more new maps and pixel positions. The GUI Import feature allows the tool to work in conjunction with coordinate outputs from libraries of automated functions and plugins from other image analysis tools. Manual selection options allow the user to interact and fine tune image and map registration. The *Project view* may be used to open image viewers of individual or aligned images and labeled pixel locations for quick assessment of alignment error (Fig. 2B, Supplementary Fig. 1D-F). When using the GUI *Wizard*, a new project can be created that will record information about maps (File), positions (Import), alignments (Align to Map), and errors that might arise during transformation (Log). New projects can be saved as XML-formatted data to preserve pre-alignment and post-alignment transformations of stage positions. The registration coordinates and corresponding stage position remain constant after transformation, allowing users to un-align and re-align the images if the transformation is unsatisfactory (Fig. 2B, Supplementary Fig. 1, Video 1). CorRelator can export multiple navigator files (Nav-2) specific to an experimental workflow and save aligned overlays for subsequent applications.

**Fig. 2.**
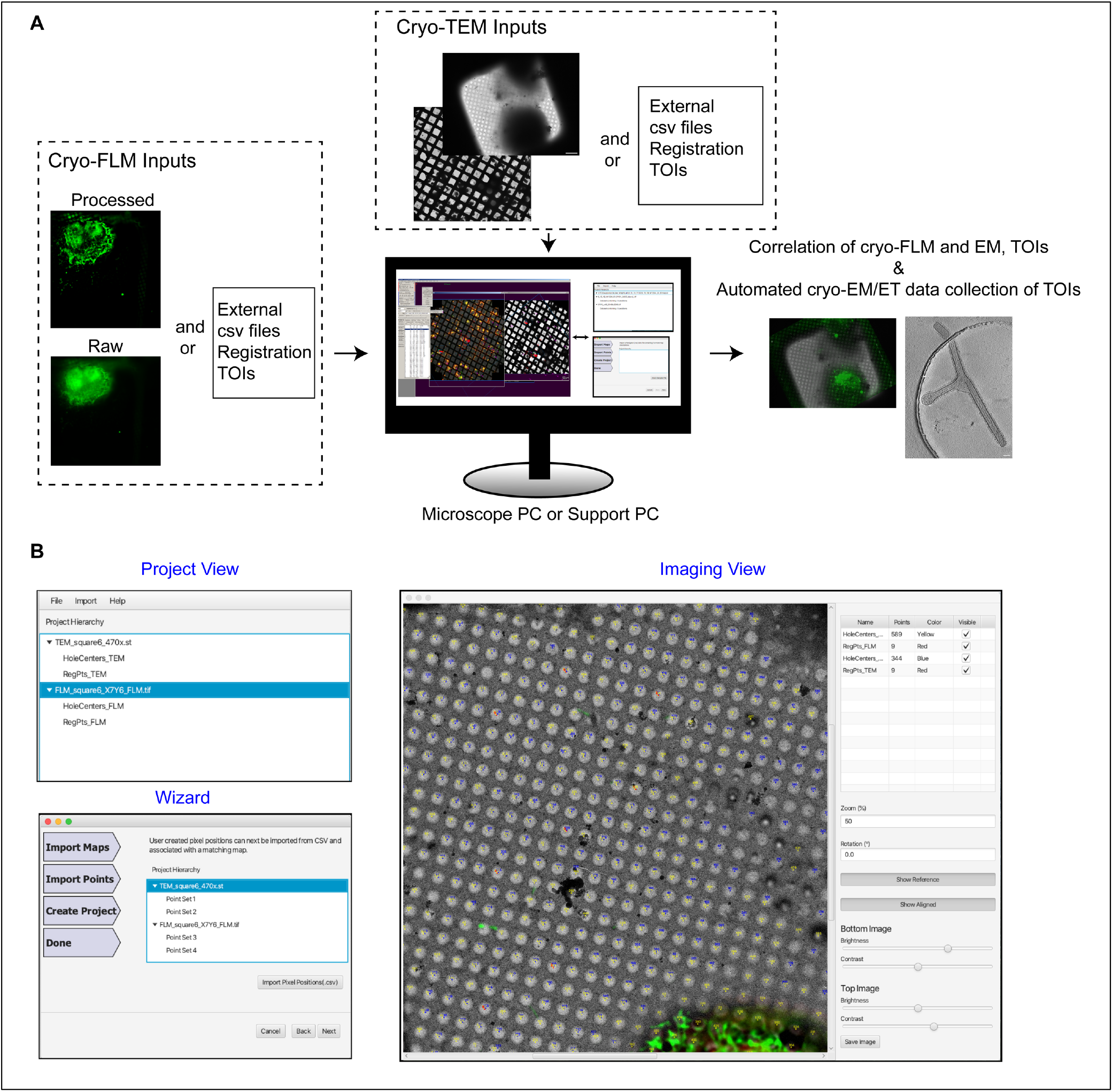
CorRelator on-the-fly cryo-CLEM workflow combined with the TEM-control program SerialEM. (A) CorRelator takes on-the-fly inputs from SerialEM including the navigator files, cryo-FLM and TEM maps and images, and optional independent csv files for registration. The resulting output can be directly imported back into SerialEM for the on-the-fly data collection of identified TOIs. (B) The CorRelator GUI features a *Project* view, an intuitive *Wizard*, and multiple image and alignment views. The GUI allows for an iterative user-in-the-loop labeling of registration positions and a quick feedback-based affine transformation to align stage positions between the FLM and TEM imaging.

### High-accuracy cryo-CLEM Applications with CorRelator

CorRelator fits into the classical two-step cryo-CLEM workflow for accurate and flexible on-the-fly correlation experiments (Fig. 3, Supplementary Fig. 2). As published previously and detailed in the Materials and Methods (Hampton et al., 2016; Ke, Dillard, et al., 2018), HeLa cells grown on a Quantifoil grid were infected by recombinant RSV strain rA2-mK^+^ (Hotard et al., 2012) expressing the far-red monomeric Katushka tag, followed by native immunogold-labeling of the RSV F glycoprotein (Fluro-nanogold) (Yi et al., 2015). We used a commercial Leica EM cryo-CLEM system and the LAS X CLEM software to scan entire grids to produce a series of multi-channel, multi-Z image stacks at each position (Hampton et al., 2016; Schorb et al., 2017). A final single stitched whole cryo-FLM grid image was then imported into SerialEM for ‘square-level’ rough correlation. Each local FLM image tile of the grid mosaic could be used later for fine correlation. Subsequent to FLM imaging, the same grid was loaded onto a Titan Krios microscope (ThermoScientific, Hillsboro, OR, USA) and a low magnification cryo-EM grid montage was collected with SerialEM for rough correlation (Supplementary Fig. 2A).

**Fig. 3.**
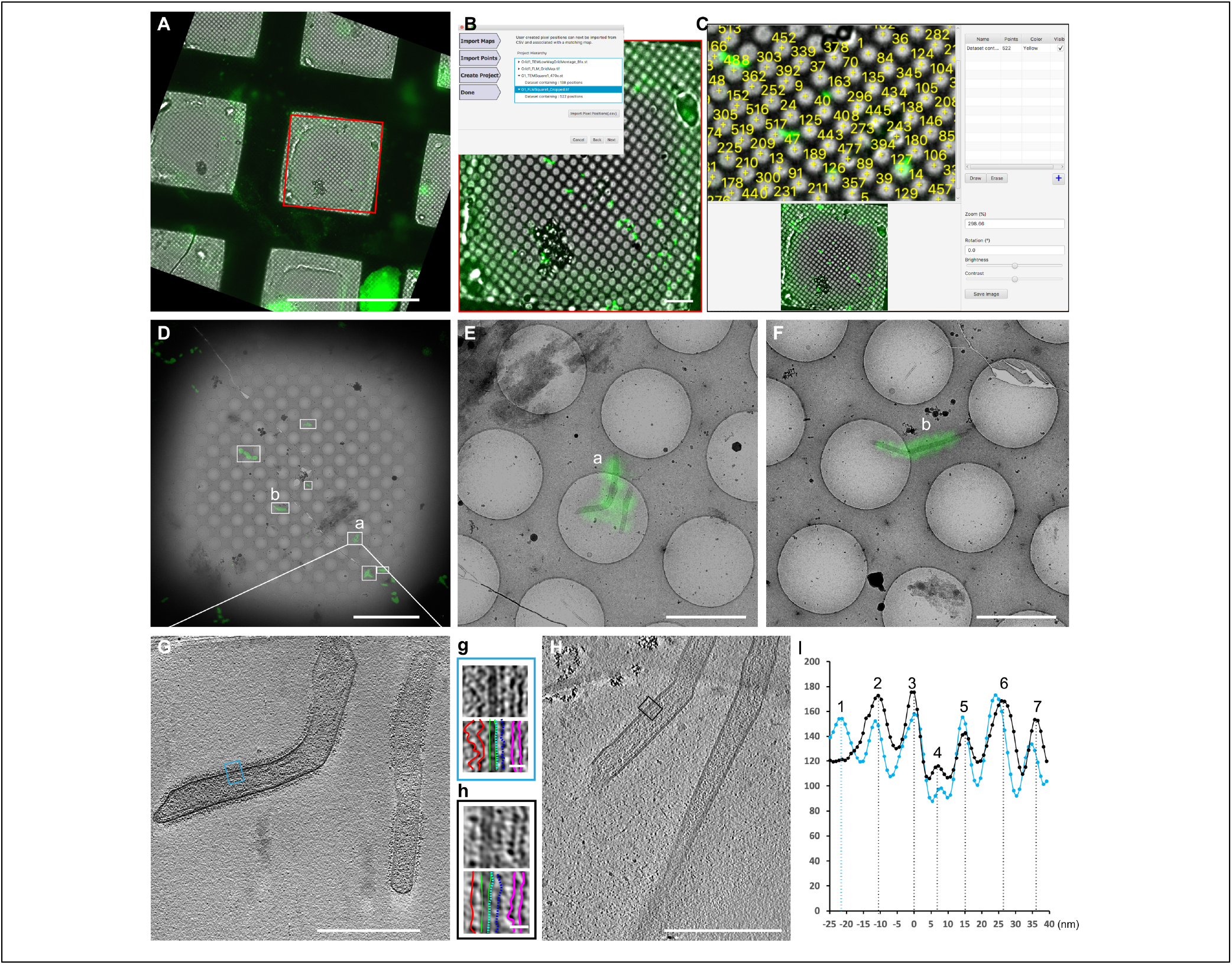
CorRelator cryo-CLEM workflow of a representative square of interest. The green fluorescence signal is indicative of Nanogold-Alexa488 immuno-labeling of the RSV F glycoprotein. (A) A raw FLM square image prior to being imported into SerialEM. The red boxed area is the cropped and rotated FLM frame imported into SerialEM, as in Supplementary Figure 2, Step 1. (B) The Wizard window that guides the user to import the SerialEM Nav_2 and external csv files into CorRelator, as in Supplementary Figure 2, Step 3. (C) The image viewer where an iterative and interactive registration is initiated for fine alignment and transformation. The imported external hole centroids (yellow) are numbered and can be visible (on and off) for guided manual registration. (D) Superposition of the correlated cryo-TEM map and −FLM map. White boxed areas indicate the TOIs where high magnification images and cryo-tilt series were recorded based on the transformed green signal. (E-F) Magnified overlays of correlated cryo-TEM and FLM mosaics of the boxed areas (a) and (b) in D. (G) A tomographic slice view (thickness of ~9 nm) corresponding to the position (a) in D and E. The inset g is a magnified view of the blue-boxed area in G. (H) A tomographic slice (thickness of ~9 nm) of released unlabeled RSV particles. The inset h is a magnified view of the black-boxed area in H. (I) The linear profiles of g (blue line) and h (black line) highlight RSV structural components: surface glycoproteins (peak 1-2, lineated in red in g and h, peak 1 corresponding to the secondary antibody nanogold-Alexa488 anti-human IgG against viral glycoproteins of peak 2), viral outer membrane (peak 3, green), viral inner membrane and a thin layer of matrix beneath (peak 4, green, cyan), M2-1 protein (peak 5, navy blue), RNP (peak 6-7, pink). Scale bars: 200 μm in A, B, 10 μm in C, D, 2 μm in E, F, 500 nm in G, H, 20 nm in g and h.

Recognizable grid-level landmarks visible in both FLM and cryo-EM such as torn and broken squares, cells, and letters or numbers (like those on an EM Finder grid), were used as rough registration points. Alignment may be difficult and time-consuming when dark regions were present in the EM image because of thick ice or when the post-FLM imaged grid was flipped or rotated during sample handling. We used the cryo-EM grid map (Supplementary Fig. 2A) as a reference to transform the raw FLM grid mosaic counterpart by flipping, rotating, and adjusting its contrast offline (Supplementary Fig. 2B, b) in Fiji, prior to importing into SerialEM. A baseline SerialEM navigator file, Nav_1, was generated to store both TEM and the imported FLM map entries. Along with Nav_1, we provided CorRelator with the independently determined coordinates in multiple csv files for registration and TOI selection (Supplementary Fig. 1). Subsequently, a Nav_2 file was generated in CorRelator and reloaded into SerialEM to display transformed fluorescent squares of interest on the TEM (Supplementary Fig. 2E-G). The rough transformation supported the identification and recording of cryo-EM frames at a medium magnification (470x, 399 Å/pixel) covering the square of interest (Supplementary Fig. 2C-E). The use of TEM images as references to guide additional image processing and registration point selection of fluorescent maps has two advantages. Frist, FLM is more tolerant to sample and ice thickness. Landmarks for registration that are visible with FLM might not be identifiable by TEM. Thus, starting with and referring to landmark selections on the TEM frames ensure an easy and accurate selection of matching registration points in both TEM and FLM images. Second, SerialEM associates acquired TEM maps to the microscope stage with a specific pixel coordinate to stage position transformation. Changes in pixel coordinates of the TEM map, especially the map’s length and width in pixels, during geometric transformation operations such as cropping, rotating, and flipping when the map’s length and width are not equivalent, could lead to the marginal misplacement of landmarks by pixels and a subsequent misalignment of the two imaging modalities. In addition, to minimize pixel changes introduced by imprecise stage shift movements, we recorded the TEM square maps at a medium low magnification (470x, EFTEM mode, Krios) where a single frame covered the entire field of view of the square of interest with the support holes clearly visible (Supplementary Fig. 2E, G, H). Thus, no additional blending or stitching of multiple image tiles was necessary.

Bead-less alignment based on sample support features visible by lower magnification imaging modes has been explored for accurate correlation (Anderson et al., 2018; Dahlberg et al., 2020). Multiple independent image analysis tools such as ImageJ/Fiji (Schindelin et al., 2012), MATLAB (MathWorks, Natick, MA, USA; (Kamarianakis et al., 2011)), SerialEM, and SerialEM with the Py-EM plugin (Schorb et al., 2019) have incorporated automated algorithmic functions to detect the edge and centroid of a hole. The hole centroid identification process may be completed in the same fluorescent channel as the TOIs, e.g., the GFP channel for labeled RSV particles, which makes the process less susceptible to misalignments and shifts between multiple frames (Schellenberger et al., 2014). Non-local means filtering can be used to increase the contrast in FLM images with high background signal (Anderson et al., 2018; Buades et al., 2005). To examine possible deviations in alignment between brightfield and fluorescence image frames, we performed the hole identification procedure on filtered fluorescence and brightfield images acquired from the same area. We found that the deviation of the hole centroid coordinates for the same hole between the two modes was tolerable (Supplementary Fig. 3E), with less than 1% of holes exceeding 2-pixel difference (399 Å/pixel). A larger number of usable center coordinates were identified in the brightfield frame (Supplementary Fig. 3D). Following the rough correlation, the local fluorescence images (Fig. 3A) were flipped, rotated, and cropped based on the TEM square counterparts prior to being imported into the Nav_2 in SerialEM. CorRelator then treated the Nav_2 as the new Nav_1 to reinitiate the alignment process (Fig. 3B). Out of the provided hole centroid coordinates from Fiji (Fig. 3C, yellow markers), a set of reference points (n = 9) were selected to calculate the matrix *M* between the two modalities. After several rounds of iterative registration and assessment, an overlay of the cryo-FLM and -EM images with the refined transformation was generated for reliable identification of the TOIs (Fig. 3D).

CorRelator mapping of TOIs from FLM to TEM can provide updated stage positions for direct automated cryo-ET tilt series collection. Here, labeled RSV particles were the targets (Fig. 3D). The positioning error of on-the-fly correlation after moving the stage to the predicted position was 218.9 nm (n = 50 targets) which was comparable to previous reports (Fu et al., 2019; Schorb et al., 2017). This positioning error was also well within targeting range for cryo-ET data collection at a higher magnification (pixel size of 3~8 Å) (Table 1). Consistent with previous reports (Hampton et al., 2016; Ke, Dillard, et al., 2018; Kiss et al., 2014; Liljeroos et al., 2013), tomographic slices of the RSV particles (Fig. 3G, H) showed that the virus was predominantly filamentous and likely infectious. Regular organization of the surface glycoproteins, matrix protein (M), M2-1, and the ribonucleoprotein complex (RNP) relative to the viral membrane were revealed in linear density profile analysis (Fig. 3G-I). Of note, the nanogold-Alexa488 labeled RSV particles appeared fuzzier than un-labeled RSV released from HeLa cells infected under the same conditions (MOI = 10, 24-hours post infection). An extra layer of density ~23 nm (peak 1) from the viral membrane was observed above the densities attributed to the glycoproteins (~12 nm, peak 2) (Fig. 3g-h, I). The extra density was likely due to the secondary antibody nanogold-Alexa488 anti-human IgG (~ 12 to 15 nm in length) and contributed to the diffuse appearance.

**Table 1.** Measurements of CorRelator target prediction performance were performed on two independent data sets of vitrified HeLa cells infected by respiratory syncytial virus (RSV) grown on two types of grids. The hole centroid pairs (n = 9) picked through automated Hough Circle Transform function in Fiji were used for registration. The non-registration hole centers (leave-out-one method) and TetraSpeck beads were considered as TOIs and used to calculate predicted errors in the application of post-acquisition deviation (left). The mean relocation error in X and Y after TEM stage movement during on-the-fly TEM relocation of vitrified RSV was calculated.

| Data Sets | Post-acquisition deviation (nm) |  |  |  | On the fly deviation (nm) |  |
| --- | --- | --- | --- | --- | --- | --- |
| | Hole Centroids<br>(2 $\mu\text{m}$) | | TetraSpeck<br>(500 nm) | | RSV particles<br>(filamentous, 1~2 $\mu\text{m}$) | |
| Sample 1: Cryo-CLEM nanogold-Alexa labeled<br>RSV on a<br>Quantifoil grid | X<br>69.8<br>(20) | Y<br>72.5<br>(20) | X<br>46.2<br>(12) | Y<br>54.2<br>(12) | X<br>147.9<br>(27) | Y<br>141.7<br>(27) |
| Sample 2: Cryo-CLEM nanogold-Alexa labeled<br>RSV on a<br>Quantifoil Finder<br>grid | 67.3<br>(24) | 61.2<br>(24) | 49.9<br>(12) | 30.8<br>(12) | 173.5<br>(23) | 155.6<br>(23) |

All map images were loaded through relative path names indicated in a navigator file. CorRelator operates on the image pixel coordinates for alignment and transformation. It is possible to use noisy unprocessed fluorescence images that have holes visible for hole-centroid registration, and then to swap with and load a processed fluorescence frame for on-the-fly target identification in SerialEM. With one consideration, there might be uncertain small changes in pixels between pre- and post-processed images. When an object is irregular and pleomorphic, such as filamentous viral particles (Fig. 3E-H, Supplementary Fig. 2I-J), locating fluorescent objects by wide-field imaging is difficult. Despite the significant advances that have been made in developing stable cryo-CLEM systems (Brandt et al., 2010; Hampton et al., 2016; Schellenberger et al., 2014; Schorb et al., 2017; Schorb & Briggs, 2014; van Driel et al., 2009), fluorescent signal from adjacent focus planes may result in an image with out-of-focus blur that limits details from being observed. To improve image contrast and resolution, we applied Leica’s THUNDER technology to remove background out-of-focus signal and enhance image contrast. An automated adaptive three-dimensional (3D) deconvolution method was applied to improve image resolution by restoring the point spread function (PSF). Here, we investigated RSV assembly sites on the plasma membrane of the infected cells (Harrison et al., 2010; Ke, Dillard, et al., 2018; Oomens et al., 2006). Labeled RSV F was resolved along the host plasma membrane and was present on the exterior of released viral particles (Ke, Dillard, et al., 2018; Oomens et al., 2006; Yi et al., 2015). After small volume computational clearance (SVCC), the image contrast and resolution were improved (Supplementary Fig. 4E). As a result, previously undistinguishable viral filaments (Fig. 4A-E) close to infected cells were observed (Fig. 4B-F). We registered and transformed unprocessed cryo-FLM (Fig. 4A) and EM images in CorRelator using paired hole centers, followed by swapping to a post-SVCC frame (Fig. 4B). SVCC processing did not introduce detectable pixel changes to the original coordinate system of the raw frame (Supplementary Fig. 4E). Correlation between the post-SVCC fluorescent image and the cryo-EM square map revealed extended viral filaments along the cell plasma membrane (Fig. 4G, Supplementary Fig. 4). Cryo-ET of the same targets (red and orange stars in Fig. 4G) highlight the spatial organization of viral components inside the RSV filament at the cell edge (Fig. 4I). The RNP is noted by white arrowheads and the F glycoprotein along the viral membrane by black arrowheads (Fig. 4I). The full-width at half-maximum (FWHM) of the PSF in the lateral X and Y directions was improved by 1.5 (n = 10), consistently with the narrower X-axial intensity distribution (red boxed signal in Supplementary Fig. 4C-D, F).

**Fig. 4.**
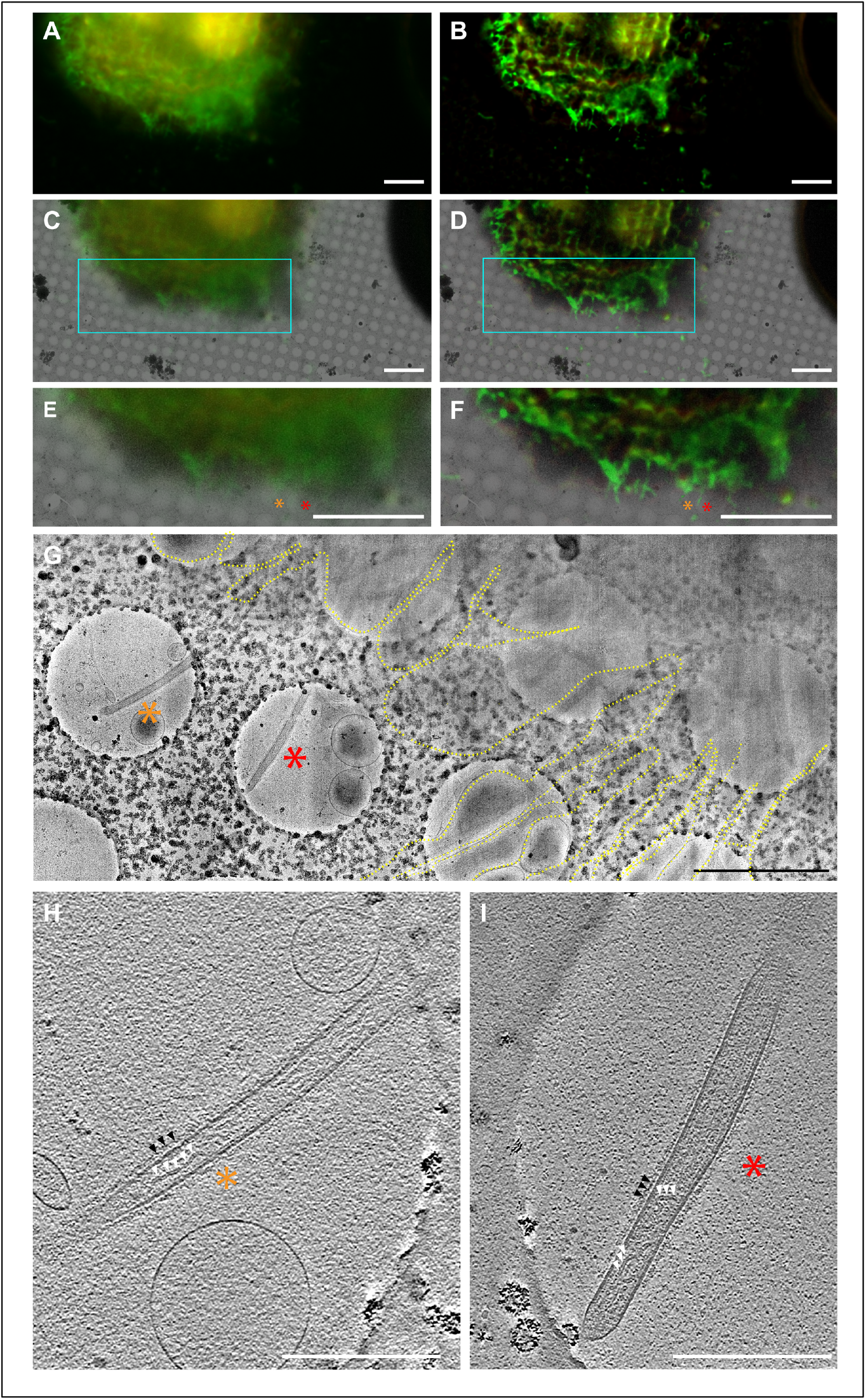
Cryo-CLEM-CorRelator with THUNDER-processed FLM images. (A) Raw widefield FLM image of an RSV-infected cell displaying a fluorescent reporter gene in RSV-infected cells (red) and labeled RSV F glycoprotein (green). (C) Overlay of a cryo-EM image and the transformed fluorescence image. (E) Magnified view of the cyan boxed area marked in C. (B, D, F) the same images A, C, E processed with THUNDER Small Volume Computational Clearance (SVCC). The orange and red asterisks indicate RSV glycoprotein fluorescent signals marked as TOIs on pre- and post-SVCC. (G) Magnified cryo-EM montage view of the star TOIs in E and F of the Nanogold-Alexa488 immuno-labeled RSV particles (green). The RSV filaments extend from the cell plasma membrane and cell protrusions (dashed yellow line). (H-I) Central sections (thickness of ~9 nm) through the tomograms collected at the marked TOIs. White triangles indicate the RSV ribonucleoprotein (RNP) inside the RSV filament. The black arrows note the RSV glycoproteins bound to antibodies and 6-nm gold (peak 1 in Fig 3I). Scale bars: 10 μm in A-F, 2 μm in G, 500 nm in H and I.

### Accuracy of correlation and error estimation

It has been reported that post-acquisition correlation in the range of 20 to 100 nm is achievable using TetraSpeck or FluoSphere fiducials combined with the MATLAB linear transformation function (Anderson et al., 2018; Kukulski et al., 2011; Paul-Gilloteaux et al., 2017; Schellenberger et al., 2014; Schorb et al., 2017). In addition, on-the-fly relocation precisions of 0.1 to 1μm may be possible when microscope stage movements are taken into consideration (Fu et al., 2019; Schorb et al., 2017). Following similar procedures (Anderson et al., 2018; Kukulski et al., 2011; Schellenberger et al., 2014), the alignment accuracy using paired hole center registration (n ≥ 7 pairs) in CorRelator was measured on full cryo-CLEM data sets. With this process, CorRelator predicted errors of transformation were between 20 and 100 nm. We found that average prediction errors in X/Y axial directions were 68.6 nm (± 42) and 66.8 nm (± 52) hole centers were treated as the objects of interest, and 48.1 nm (± 43) and 42.8 nm (± 34) for 500 nm TetraSpeck targets (Table 1, Supplementary Fig. 5A). The slight improvement seen in TetraSpeck prediction, was likely due to the application of a 2D Gaussian fitting model to detect precise coordinates of the point-like fluorescent signal (Kukulski et al., 2011; Schellenberger et al., 2014). Next, we tested the on-the-fly target prediction on pleomorphic virus particles. To demonstrate the image acquisition procedure, we applied a similar methodology (Schorb et al., 2017) and analyzed a set of 50 images acquired at magnifications of 2,250x (76 Å/pixel) and 19,500x (4.47 Å/pixel). After performing the transformation with 470x EM images, we moved to the stage positions of the transformed fluorescent signal and recorded images at 2,250x and 19,500x. For best results in SerialEM, image shift calibrations should be performed between the registration magnification (470x), medium magnification (2,250x), and acquisition magnification (19,500x). Taking into account variations in stage movement, the Euclidian distance between the actual position of the fluorescent viral particle (pink cross, Fig. 5C-G) and the center of the images (or predicted position, yellow cross, Fig. 5C-G) was used to estimate the on-the-fly alignment accuracy, as described previously (Schorb et al., 2017). The mean error of Euclidian distance was 218.9 nm with a standard deviation of 109 nm, with an average of 160.7 nm in X and 148.6 nm in Y axis (n = 50, Fig. 6L, Table 1). A maximum prediction error of 605 nm was observed when the hole centroids over damaged support films were used for image registration (Fig. 5F). The error for positioning pleomorphic objects fell well within the cryo-EM imaging tolerance range of ~1.4 x 1.4 μm at a pixel size of 4.47 Å on a Gatan K3 direct electron detector. The approximate normal distribution of prediction errors indicated that data acquisition could be conducted with a field of view as small as 700 nm while still expecting a < 5% off-target rate (Fig. 6L). We noted that on-the-fly accuracy depended on multiple factors, including registration point selection, stage and image calibrations between magnifications on the TEM, and the intensity and resolution of the fluorescent signals in FLM images. We obtained consistent targeting precision when stage and image shifts were calibrated at the magnifications used for correlation in SerialEM (Mastronarde, 2005).

**Fig. 5.**
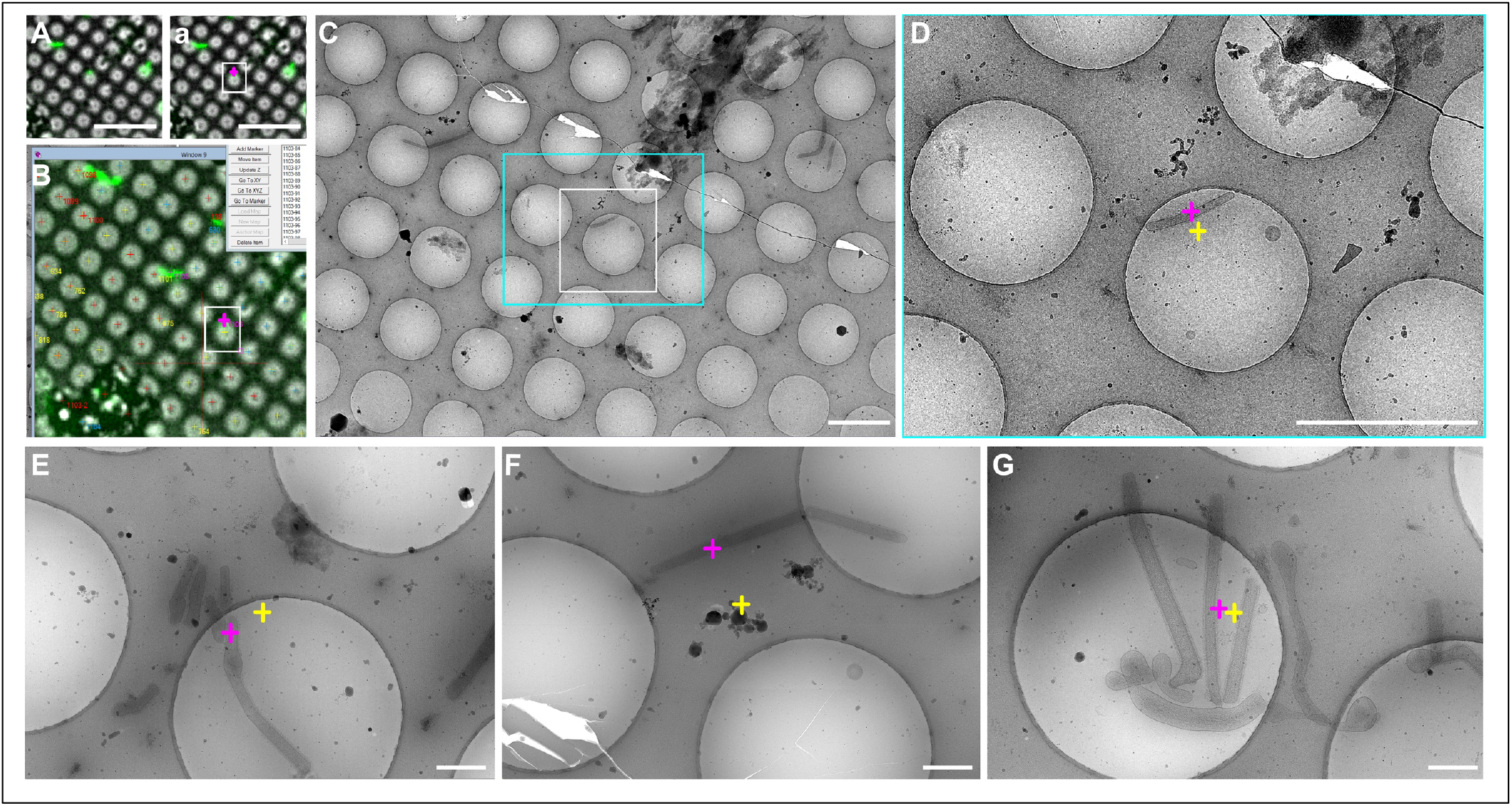
Correlation precision of on-the-fly Cryo-CLEM-CorRelator for labeled RSV particles. (A) Single bright-field and fluorescent channel merged FLM map used to provide the TOIs (white boxed area in (a). The labeled RSV filaments appear green. (B) SerialEM screenshot of the post-correlated FLM map of the same region in (A) and white boxed area in (a). The top right is part of the Nav_2 written out by CorRelator after being reloaded in SerialEM. The pink point indicates the center of the virus (green) as a representative TOI in the post-correlated FLM map. (C) Higher magnification cryo-EM image of the same white boxed hole in (a) and (B) and its surrounding area, after moving the TEM stage to the TOI (pink cross in (a), (B)). (D) Zoomed view into the cyan boxed area in (C). The center coordinate (yellow cross in (D-G)) of each TEM frame corresponds to the predicted stage position by CorRelator after moving the stage. The pink cross in (D-G) indicates the actual TOIs. (E-G) High magnification TEM images that are analogous coordinates for both actual (pink cross) and predicted positions (yellow cross). Scale bars: 10 μm in A and a, 2 μm in C-D, 500 nm in E-G.

**Fig. 6.**
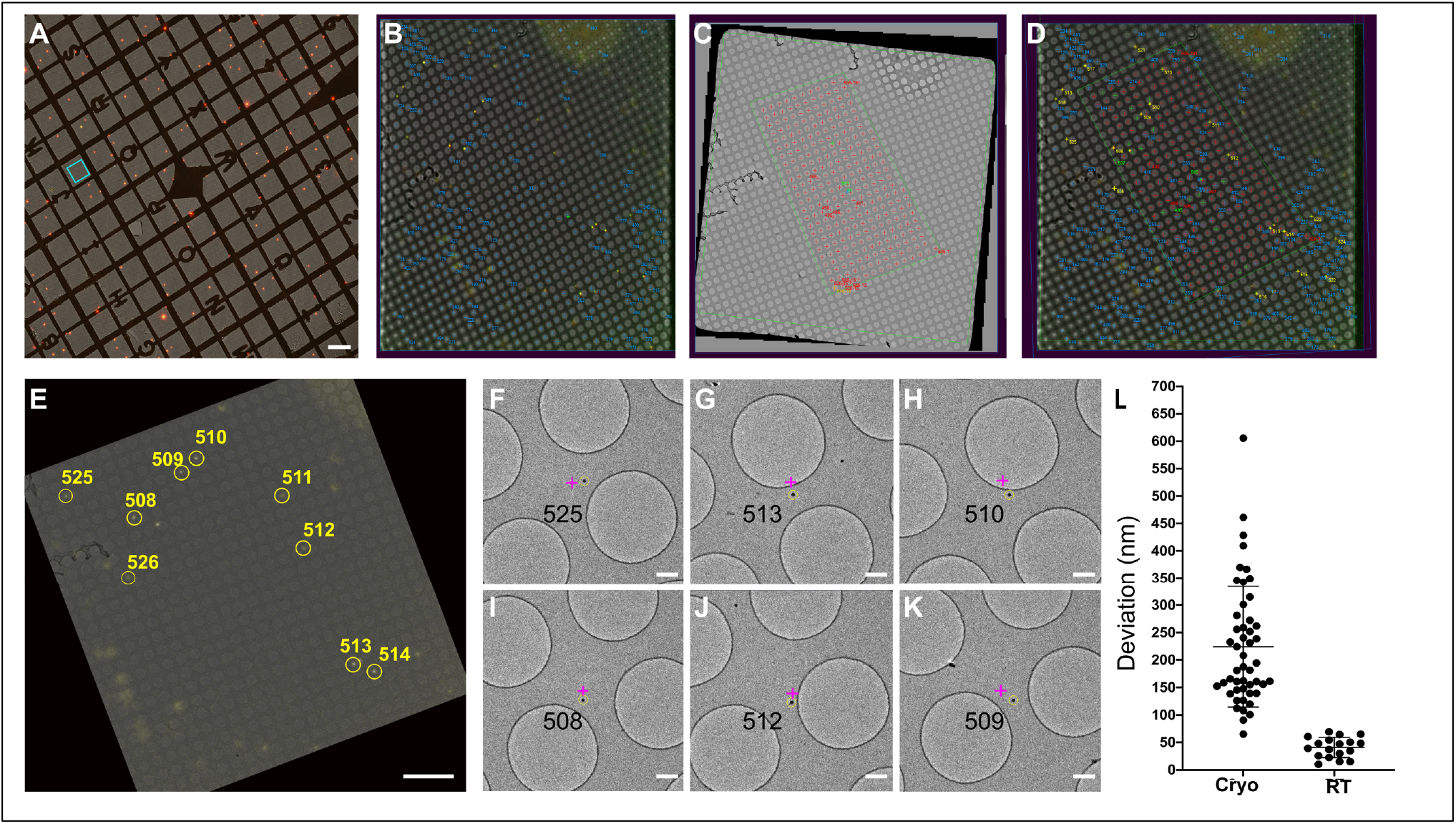
Correlation of TetraSpeck beads under ambient conditions (room temperature). (A) FLM grid montage of a TetraSpeck-coated Quantifoil Finder grid. (B) SerialEM screenshot of a single FLM image of the region corresponding to the cyan boxed area in (A). Hole centers were identified offline with Fiji (blue crosses) and displayed as imported “external” stage positions after reloading the Nav_2 written by CorRelator, prior to correlation in SerialEM. (C) SerialEM Screenshot of a single medium-magnification TEM image corresponding to the square image of (B). Hole centers (red cross) were identified online with SerialEM ‘Add Grid of Points’ function to fill a polygon (green) item. (D) Post-correlation screenshot of the FLM map (B) and TEM map (C) in SerialEM. The hole center stage positions (C) (red) were transferred into the FLM map after correlation. The hole centers that were used for registration are marked in green, while numbered yellow points (508-524) were marked as the TOIs. (E) Superposition of the TEM and FLM correlated maps. The same TOIs in D were circled in yellow. (F-G) Acquired high-magnification images at the yellow TOIs in D and E after moving the stage to the predicted positions (pink cross). The yellow circle was centered on the actual TOIs using the 2D Gaussian fit. The pink cross marks the predicted coordinates of the TOIs calculated by CorRelator. (L) Distribution plot of the coordinate deviation by CorRelator between the actual TOIs and predicated positions under cryogenic (n = 50) and ambient conditions (n = 19). TOIs are TetraSpeck beads for ambient CLEM. Scale bars: 200 μm in A, 10 μm in E, 500 nm in F-G.

We also compared the transformation of CorRelator to standard MATLAB affine and projective geometric functions that have been used for consistent high accuracy correlation (Fu et al., 2019; Kukulski et al., 2011; Schellenberger et al., 2014; Schorb et al., 2017) and eC-CLEM (Paul-Gilloteaux et al., 2017). For comparison, the identical set of paired hole centroids (n = 7 to 10) on cryo-FLM and cryo-EM images were used for registration while the prediction errors were measured on the same coordinates for fluorescent RSV particles. CorRelator performed equivalently to the MATLAB Affine Transformation and eC-CLEM Rigid Transformation that corrects translation, rotation, scaling, and shearing, the performance of which was comparable to, and slightly better than, MATLAB’s Projective Transformation function (Supplementary Fig. 5B, Fig. 7B; (Paul-Gilloteaux et al., 2017)). Application of the coordinate transform from cryo-FLM to TEM showed excellent overlap between hole centroids (Supplementary Fig. 5C-D). We demonstrated that CorRelator can handle moderate grid bending that may be introduced during the handling of cryo-grids (Supplementary Fig. 2).

### Adaptability of the workflow

To assess CorRelator integration with room temperature FLM-EM correlative imaging schemes, we used TetraSpeck-coated EM Finder grids and conducted room-temperature FLM-EM correlations. We designed this prototype application to determine the: (1) potential adaptability of CorRelator for customized applications; and (2) alignment accuracy through hole center registration in the absence of cryogenic handling and thermal stage drift observed with cryo-fluorescence microscopy systems.

The number of 100 nm TetraSpeck beads was optimized to roughly 5~15 beads per square of an EM Finder grid (Schellenberger et al., 2014). A multi-channel automated scan of the whole grid was performed by Micro-Manager (Edelstein et al., 2010) on an inverted DMi8 microscope at 40x (40 x, 0.6 NA, dry, Leica). The scanned tiles were stitched together to generate a single FLM specimen frame for grid-wide rough correlation (Fig. 6A, Supplementary Fig. 2 Step 1). A higher NA oil-immersion objective (63 x, 1.4 NA, oil-immersion, Leica) was used to record single square frames of areas with a good bead distribution. The GFP channel square frame either went through hole center identification (Fig. 6B) or a 2D Gaussian fitting to determine precise bead coordinates (Schellenberger et al., 2014). The same grid was then loaded onto a Tecnai T12 microscope (ThermoScientific, Hillsboro, OR, USA) and a whole grid montage and associated navigator file (Nav_1) were written out by SerialEM. The rough correlation on the grid-level was done as described above (Supplementary Fig. 2 Step 1), followed by the TEM square image collection.

For fine correlation (Supplementary Fig. 2 Step 3), we used the hole center coordinates of FLM and EM, that were externally identified with Hough Circle Transform in Fiji, and refined by the user prior to being transformed into stage positions in CorRelator (Route 2 in Fig. 1), referred to as Approach 1. We could also directly mark the hole centers as stage positions on the TEM (not applicable to the FLM image) using the SerialEM ‘Add Grid of Points’ function (Fig. 6C) based on the unguided manual lattice pattern prediction. We referred to this hybrid point-selection approach as Approach 2. In both cases, one-to-one hole-center based registration between the FLM and EM was determined, followed by a transformation in SerialEM using its ‘Transform’ function. We verified the success of the on-the-fly correlation by analyzing the acquired image sets after moving the stage to the predicted positions of the 100 nm beads treated as the TOIs here (Fig. 6). In Approach 1, the mean displacement of the beads was 40.8 nm (n = 19, S.D. = 18 nm, Fig. 6F-L, Table 2), while the Euclidian deviation of the positions of the beads from the center of the images was 120 nm (n = 19, S.D. = 32 nm, Table 2) in the hybrid Approach 2. The smaller prediction errors in the first scenario suggested the practical necessity of combining automated algorithmic approaches and user interaction. Overall, we show that CorRelator may be adapted for use with many microscope systems and is sufficiently flexible to be incorporated into common SerialEM operations to achieve consistent high accuracy correlation.

**Table 2.** Measurements of CorRelator target prediction performance were performed on a TetraSpeck-coated Finder grid imaged at ambient condition. On the left: the hole centroid pairs (n = 9) picked through automated Hough Circle Transform function in Fiji were used for registration. The non-registration hole centers (leave-out-one method) and TetraSpeck beads were considered as TOIs and used to calculate predicted errors in the application of post-acquisition deviation. On the right: The mean relocation error in X and Y after TEM stage movement during on-the-fly TEM correlation of TetraSpeck targets was calculated. The functions ‘Add Grid of Points’ and ‘Transform Items’ in SerialEM were applied to identify hole centroids on the TEM map and to correlate two modalities. SerialEM column: the hole identification of FLM was done in CorRelator while the hole identification of TEM, registration, and transformation of both modalities were performed in SerialEM. CorRelator column: the hole identification, registration, and transformation of FLM and TEM were both done in CorRelator. The output Nav_2 was then reloaded in SerialEM.

| Data Sets | Post-acquisition deviation (nm) |  |  |  | On the fly deviation (nm) |  |  |  |
| --- | --- | --- | --- | --- | --- | --- | --- | --- |
| | Hole Centroids<br>(2 μm) | | TetraSpeck<br>(100nm) | | SerialEM | | CorRelator | |
| Sample 3:<br>FluoSphere of<br>100 nm on a<br>Quantifoil<br>Finder grid at<br>room<br>temperature | X | Y | X | Y | X | Y | X | Y |
|  | 14.1<br>(12) | 16.7<br>(12) | 4.8<br>(12) | 5.7<br>(12) | 92.2<br>(19) | 76.7<br>(19) | 28.4<br>(19) | 29.0<br>(19) |

## Discussion

The diversity of FLM and cryo-FLM modalities in combination with TEM has led to advances in workflow and software development (Supplementary Table 1). Here, we introduce a new flexible CLEM application tool, CorRelator (Fig. 1, Supplementary Fig. 6), that can be coupled with SerialEM for TEM data collection. We developed CorRelator to facilitate the integration of high-precision target identification by FLM with automated high-throughput cryo-EM/ET data collection. Using the interactive hole centroids-based alignment, we demonstrate that CorRelator is able to achieve and improve the standard bead-aided prediction error of 20 to 100 nm for 2D post-acquisition cryo-correlation, and 100 to 700 nm relocation precision for on-the-fly cryo-EM/ET acquisition. We explored the use of advanced image processing steps in regular cryo-CLEM workflows, such as computational cleaning and 3D deconvolution approaches. Due to its simplicity, we show that CorRelator could be adapted to any existing FLM, cryo-FLM, or TEM system, expanding the scope of correlative microscopy. CorRelator aims to bridge between cryo-FLM and on-the-fly cryo-EM for an easy and accurate transition. This means that the main tasks in building a correlative imaging pipeline to support automated cryo-EM/ET data collection is now reduced to user-specific experimental designs and unambiguous TOI identification.

To achieve high-accuracy correlation with CorRelator, no specialized hardware requirements are needed for the FLM and TEM microscopes and cameras. Good microscope column alignment and camera calibrations ensure a good, nonradical transformation estimation from 3-D to 2-D, from the real object space (to camera coordinates) to image (film) space, then to pixel coordinates displayed on the screen (Gonzalez & Woods, 2002; Szeliski, 2011). Good SerialEM calibrations on the TEM that support Navigator usage makes stage movement and image acquisition more robust. Experiments were performed with a Titan Krios, Tecnai TF30 (data not shown), Tecnai T12, and Talos L120C (data not shown). CorRelator’s transformation performance has not been tested on plastic-embedded or high-pressure frozen/freeze substituted samples (Kukulski et al., 2011; Paul-Gilloteaux et al., 2017) where larger structural changes and sample deformations can occur that may require dedicated transformation parameter selection (Paul-Gilloteaux et al., 2017).

We compared the 2D transformation performance of CorRelator to the registration/correlation functions in MATLAB (Fu et al., 2019; Kukulski et al., 2011; Schellenberger et al., 2014; Schorb et al., 2017) and eC-CLEM (Paul-Gilloteaux et al., 2017) (Supplementary Fig. 5B, 7B). We adopted the linear closed-form ordinary least square (OLS) solution to an overdetermined system (Horn, 1987). The use of closed-form least square normal method has two main advantages. It is preferred over manual and semi-automated point-matching registration where a “smaller” set of paired reference points are usually provided, speeding up the matrix computing and calculation. Second, it limits unnecessary liberty in a non-rigid, non-linear transform that could warp and distort images that lead to increased target registration error in correlation (Supplementary Fig. 5B, Fig. 7B; Paul-Gilloteaux et al., 2017).

The closed-form solution and non-linear product transformation require at least four reference points to avoid a singular/coplanar matrix, as is always the case when there are only three reference coordinates between two coordinate systems (Gonzalez & Woods, 2002; Horn, 1987). To validate the variance in transformation performance caused by selection of alignment markers, we used a set of five reference coordinate pairs of hole centroids, defined either locally within a bounding area of ~3% of the entire field of view, or across the entire image spanning roughly 50%, to obtain the transformation matrices in CorRelator, MATLAB (affine and projective transformation), and eC-CLEM (rigid transformation). The calculated matrices were then applied to the same set of TOIs (leave-out-one, non-registration hole centroids) to obtain the prediction errors. We found that n ≥ 5 reference coordinate pairs were able to achieve an overall high-accuracy cryo-FLM-to-TEM coordinate correlation when the registration pairs were roughly distributed across 50% of the entire image (Supplementary Fig. 5D). The prediction error for TOIs that were closer to the picked reference coordinates were smaller than those farther away. Increasing the number of local reference coordinates marginally improved the prediction error. We note that more uniformly distributed reference coordinates help obtain an optimum cryo-FLM-to-TEM correlation. Compared to fluorescent bead-aided alignment, the distribution of holes is regular, uniform, and abundant by nature, more tolerable to prediction error caused by mis-position of reference pairs.

There is an increasing need for fast, accurate, and automated correlative registration for higher throughput. We validated the potential improvements that CorRelator can provide in robust alignments using highly interactive workflows. After importing algorithmically-determined external coordinates, the GUI-driven manual registration and image viewing assessment supports users during registration point reassignment and editing. This feature is essential for areas that contain many TOIs but may lack registration accuracy. A recent report on cryo-super resolution CLEM has shown that the most robust and accurate registrations are from manual hole center identifications, as opposed to purely algorithmic approaches (Dahlberg et al., 2020). CorRelator directly supports iterative and manual selections. Further developments in CorRelator to extend and enhance its capabilities will include: image analysis tools and workflows to support cryo-FLM-FIB-milling (Arnold et al., 2016; Gorelick et al., 2019; Hsieh et al., 2014; Michael Marko et al., 2007; M. Marko et al., 2006; Rigort et al., 2010; Zachs et al., 2020).

## Acknowledgements

We thank Dr. Larry J. Anderson and Dr. Binh Ha in the Department of Pediatrics, Emory University for generously sharing motavizumab (primary antibody against respiratory syncytial virus glycoprotein F). We thank Dr. Louise Bertrand and Dr. Vikram Kohli for sharing light microscopy and cryo-FLM experience and supporting the Leica THUNDER SVCC hardware. This work was supported in part by the University of Wisconsin, Madison, the Department of Biochemistry at the University of Wisconsin, Madison, and public health service grants R01 GM114561, R01 GM104540, and R01 GM104540-03W1 to E.R.W. from the NIH. The authors gratefully acknowledge use of facilities and instrumentation at the UW-Madison Wisconsin Centers for Nanoscale Technology (wcnt.wisc.edu), which is partially supported by the NSF through the University of Wisconsin Materials Research Science and Engineering Center (DMR-1720415). A portion of this research was supported by NIH grant U24GM129547 and performed at the PNCC at OHSU and accessed through EMSL (grid.436923.9), a DOE Office of Science User Facility sponsored by the Office of Biological and Environmental Research. We are also grateful for the use of facilities and instrumentation at the Cryo-EM Research Center in the Department of Biochemistry at the University of Wisconsin, Madison during CorRelator’s performance optimization.

## References

Agronskaia, A. V., Valentijn, J. A., van Driel, L. F., Schneijdenberg, C. T. W. M., Humbel, B. M., van Bergen en Henegouwen, P. M. P., Verkleij, A. J., Koster, A. J., & Gerritsen, H. C. (2008). Integrated fluorescence and transmission electron microscopy. Journal of Structural Biology, 164(2) 183–189. https://doi.org/10.1016/j.jsb.2008.07.003

Andersen, M., Barker, B., Chou, A., Diggory, M., Donkin, R., O’Brien, T., Maisonobe, L., Pietschmann, J., Pourbaix, D., Steitz, P., & Worden, B. (n.d.). Commons math: The apache commons mathematics library. Retrieved August 1, 2016, from http://commons.apache.org/math/, online. 2011

Anderson, K. L., Page, C., Swift, M. F., Hanein, D., & Volkmann, N. (2018). Marker-free method for accurate alignment between correlated light, cryo-light, and electron cryo-microscopy data using sample support features. Journal of Structural Biology, 201(1), 46–51. https://doi.org/10.1016/j.jsb.2017.11.001

Arnold, J., Mahamid, J., Lucic, V., de Marco, A., Fernandez, J.-J., Laugks, T., Mayer, T., Hyman, A. A., Baumeister, W., & Plitzko, J. M. (2016). Site-Specific Cryo-focused Ion Beam Sample Preparation Guided by 3D Correlative Microscopy. Biophysical Journal, 110(4), 860–869. https://doi.org/10.1016/j.bpj.2015.10.053

Bharat, T. A. M., Kureisaite-Ciziene, D., Hardy, G. G., Yu, E. W., Devant, J. M., Hagen, W. J. H., Brun, Y. V., Briggs, J. A. G., & Löwe, J. (2017). Structure of the hexagonal surface layer on Caulobacter crescentus cells. Nature Microbiology, 2(7) 17059. https://doi.org/10.1038/nmicrobiol.2017.59

Brandt, F., Carlson, L.-A., Hartl, F. U., Baumeister, W., & Grünewald, K. (2010). The Three-Dimensional Organization of Polyribosomes in Intact Human Cells. Molecular Cell, 39(4), 560–569. https://doi.org/10.1016/j.molcel.2010.08.003

Briegel, A., Chen, S., Koster, A. J., Plitzko, J. M., Schwartz, C. L., & Jensen, G. J. (2010). Correlated Light and Electron Cryo-Microscopy. In Methods in Enzymology (Vol. 481, pp. 317–341). Elsevier. https://doi.org/10.1016/S0076-6879(10)81013-4

Briggs, J. A. (2013). Structural biology in situ—The potential of subtomogram averaging. Current Opinion in Structural Biology, 23(2), 261–267. https://doi.org/10.1016/j.sbi.2013.02.003

Buades, A., Coll, B., & Morel, J.-M. (2005). A Non-Local Algorithm for Image Denoising. 2005 IEEE Computer Society Conference on Computer Vision and Pattern Recognition (CVPR’05). 2, 60–65. https://doi.org/10.1109/CVPR.2005.38

Canny, J. (1986). A computational approach to edge detection. IEEE Transactions on Pattern Analysis and Machine Intelligence, 8(6), 679–698.

Castleman, K. R. (1995). Digital Image Processing (Third).

Chang, Y.-W., Chen, S., Tocheva, E. I., Treuner-Lange, A., Löbach, S., Søgaard-Andersen, L., & Jensen, G. J. (2014). Correlated cryogenic photoactivated localization microscopy and cryo-electron tomography. Nature Methods, 11(?), 737–739. https://doi.org/10.1038/nmeth.2961

Cheutin, T., Sauvage, C., Tchélidzé, P., O’Donohue, M. F., Kaplan, H., Beorchia, A., & Ploton, D. (2007). Visualizing Macromolecules with Fluoronanogold: From Photon Microscopy to Electron Tomography. In Methods in Cell Biology (Vol. 79, pp. 559–574). Elsevier. https://doi.org/10.1016/S0091-679X(06)79022-7

Dahlberg, P. D., Saurabh, S., Sartor, A. M., Wang, J., Mitchell, P. G., Chiu, W., Shapiro, L., & Moerner, W. E. (2020). Cryogenic single-molecule fluorescence annotations for electron tomography reveal in situ organization of key proteins in *Caulobacter*. Proceedings of the National Academy of Sciences, 202001849. https://doi.org/10.1073/pnas.2001849117

Dick, R. A., Xu, C., Morado, D. R., Kravchuk, V., Ricana, C. L., Lyddon, T. D., Broad, A. M., Feathers, J. R., Johnson, M. C., Vogt, V. M., Perilla, J. R., Briggs, J. A. G., & Schur, F. K. M. (2020). Structures of immature EIAV Gag lattices reveal a conserved role for IP6 in lentivirus assembly. PLOS Pathogens, 16(1), e1008277. https://doi.org/10.1371/journal.ppat.1008277

Edelstein, A., Amodaj, N., Hoover, K., Vale, R., & Stuurman, N. (2010). Computer Control of Microscopes Using μManager. Current Protocols in Molecular Biology, 92(1). https://doi.org/10.1002/0471142727.mb1420s92

Edelstein, A., Tsuchida, M. A., Amodaj, N., Pinkard, H., Vale, R. D., & Stuurman, N. (2014). Advanced methods of microscope control using μManager software. Journal of Biological Methods, 1(2) 10. https://doi.org/10.14440/jbm.2014.36

Erlendsson, S., Morado, D. R., Cullen, H. B., Feschotte, C., Shepherd, J. D., & Briggs, J. A. G. (2020). Structures of virus-like capsids formed by the Drosophila neuronal Arc proteins. Nature Neuroscience, 23(2), 172–175. https://doi.org/10.1038/s41593-019-0569-y

Faas, F. G. A., Bárcena, M., Agronskaia, A. V., Gerritsen, H. C., Moscicka, K. B., Diebolder, C. A., van Driel, L. F., Limpens, R. W. A. L., Bos, E., Ravelli, R. B. G., Koning, R. I., & Koster, A. J. (2013). Localization of fluorescently labeled structures in frozen-hydrated samples using integrated light electron microscopy. Journal of Structural Biology, 181(3), 283–290. https://doi.org/10.1016/j.jsb.2012.12.004

Fu, X., Ning, J., Zhong, Z., Ambrose, Z., Charles Watkins, S., & Zhang, P. (2019). AutoCLEM: An Automated Workflow for Correlative Live-Cell Fluorescence Microscopy and Cryo-Electron Tomography. Scientific Reports, 9(1), 19207. https://doi.org/10.1038/s41598-019-55766-8

Gonzalez, R. C., & Woods, R. E. (2002). Digital Image Processing (Second). Prentice Hall.

Gorelick, S., Buckley, G., Gervinskas, G., Johnson, T. K., Handley, A., Caggiano, M. P., Whisstock, J. C., Pocock, R., & de Marco, A. (2019). PIE-scope, integrated cryo-correlative light and FIB/SEM microscopy. ELife, 8, e45919. https://doi.org/10.7554/eLife.45919

Hampton, C. M., Strauss, J. D., Ke, Z., Dillard, R. S., Hammonds, J. E., Alonas, E., Desai, T. M., Marin, M., Storms, R. E., Leon, F., Melikyan, G. B., Santangelo, P. J., Spearman, P. W., & Wright, E. R. (2016). Correlated fluorescence microscopy and cryo-electron tomography of virus-infected or transfected mammalian cells. Nature Protocols, 12(1), 150–167. https://doi.org/10.1038/nprot.2016.168

Harrison, M. S., Sakaguchi, T., & Schmitt, A. P. (2010). Paramyxovirus assembly and budding: Building particles that transmit infections. The International Journal of Biochemistry & Cell Biology, 42(9), 1416–1429. https://doi.org/10.1016/j.biocel.2010.04.005

Horn, B. K. P. (1987). Closed-form solution of absolute orientation using unit quaternions. Journal of the Optical Society of America A, 4(4), 629. https://doi.org/10.1364/JOSAA.4.000629

Hotard, A. L., Shaikh, F. Y., Lee, S., Yan, D., Teng, M. N., Plemper, R. K., Crowe, J. E., & Moore, M. L. (2012). A stabilized respiratory syncytial virus reverse genetics system amenable to recombination-mediated mutagenesis. Virology, 434(1), 129–136. https://doi.org/10.1016/j.virol.2012.09.022

Hsieh, C., Schmelzer, T., Kishchenko, G., Wagenknecht, T., & Marko, M. (2014). Practical workflow for cryo focused-ion-beam milling of tissues and cells for cryo-TEM tomography. Journal of Structural Biology, 185(1), 32–41. https://doi.org/10.1016/j.jsb.2013.10.019

Illingworth, J., & Kittler, J. (1987). The Adaptive Hough Transform. IEEE Transactions on Pattern Analysis and Machine Intelligence, PAMI-9(5), 690–698. https://doi.org/10.1109/TPAMI.1987.4767964

Jun, S., Ke, D., Debiec, K., Zhao, G., Meng, X., Ambrose, Z., Gibson, G. A., Watkins, S. C., & Zhang, P. (2011). Direct Visualization of HIV-1 with Correlative Live-Cell Microscopy and Cryo-Electron Tomography. Structure, 19(11), 1573–1581. https://doi.org/10.1016/j.str.2011.09.006

Jun, S., Ro, H.-J., Bharda, A., Kim, S. I., Jeoung, D., & Jung, H. S. (2019). Advances in Cryo-Correlative Light and Electron Microscopy: Applications for Studying Molecular and Cellular Events. The Protein Journal, 38(6), 609–615. https://doi.org/10.1007/s10930-019-09856-1

Kamarianakis, Z., Buliev, I., & Pallikarakis, N. (2011). Robust identification and localization of intramedullary nail holes for distal locking using CBCT: A simulation study. Medical Engineering & Physics, 33(4), 479–489. https://doi.org/10.1016/j.medengphy.2010.11.016

Kaufmann, R., Schellenberger, P., Seiradake, E., Dobbie, I. M., Jones, E. Y., Davis, I., Hagen, C., & Grünewald, K. (2014). Super-Resolution Microscopy Using Standard Fluorescent Proteins in Intact Cells under Cryo-Conditions. Nano Letters, 14(7), 4171–4175. https://doi.org/10.1021/nl501870p

Ke, Z., Dillard, R. S., Chirkova, T., Leon, F., Stobart, C. C., Hampton, C. M., Strauss, J. D., Rajan, D., Rostad, C. A., Taylor, J. V., Yi, H., Shah, R., Jin, M., Hartert, T. V., Peebles, R. S., Graham, B. S., Moore, M. L., Anderson, L. J., & Wright, E. R. (2018). The Morphology and Assembly of Respiratory Syncytial Virus Revealed by Cryo-Electron Tomography. Viruses, 10(8). https://doi.org/10.3390/v10080446

Ke, Z., Strauss, J. D., Hampton, C. M., Brindley, M. A., Dillard, R. S., Leon, F., Lamb, K. M., Plemper, R. K., & Wright, E. R. (2018). Promotion of virus assembly and organization by the measles virus matrix protein. Nature Communications, 9(1), 1736. https://doi.org/10.1038/s41467-018-04058-2

Kiss, G., Holl, J. M., Williams, G. M., Alonas, E., Vanover, D., Lifland, A. W., Gudheti, M., Guerrero-Ferreira, R. C., Nair, V., Yi, H., Graham, B. S., Santangelo, P. J., & Wright, E. R. (2014). Structural Analysis of Respiratory Syncytial Virus Reveals the Position of M2-1 between the Matrix Protein and the Ribonucleoprotein Complex. Journal of Virology, 88(13), 7602–7617. https://doi.org/10.1128/JVI.00256-14

Koning, R. I., Celler, K., Willemse, J., Bos, E., van Wezel, G. P., & Koster, A. J. (2014). Correlative Cryo-Fluorescence Light Microscopy and Cryo-Electron Tomography of Streptomyces. In Methods in Cell Biology (Vol. 124, pp. 217–239). Elsevier. https://doi.org/10.1016/B978-0-12-801075-4.00010-0

Koster, A. J., & Grünewald, K. (2014). Editorial on Correlative microscopy. Ultramicroscopy, 143, 1–2. https://doi.org/10.1016/j.ultramic.2014.03.010

Kovtun, O., Leneva, N., Bykov, Y. S., Ariotti, N., Teasdale, R. D., Schaffer, M., Engel, B. D., Owen, David. J., Briggs, J. A. G., & Collins, B. M. (2018). Structure of the membrane-assembled retromer coat determined by cryo-electron tomography. Nature, 561(7724), 561–564. https://doi.org/10.1038/s41586-018-0526-z

Kremer, J. R., Mastronarde, D. N., & McIntosh, J. R. (1996). Computer visualization of three-dimensional image data using IMOD. Journal of Structural Biology, 116(1), 71–76. https://doi.org/10.1006/jsbi.1996.0013

Kukulski, W., Schorb, M., Welsch, S., Picco, A., Kaksonen, M., & Briggs, J. A. G. (2011). Correlated fluorescence and 3D electron microscopy with high sensitivity and spatial precision. The Journal of Cell Biology, 192(1), 111–119. https://doi.org/10.1083/jcb.201009037

Li, S., Ji, G., Shi, Y., Klausen, L. H., Niu, T., Wang, S., Huang, X., Ding, W., Zhang, X., Dong, M., Xu, W., & Sun, F. (2018). High-vacuum optical platform for cryo-CLEM (HOPE): A new solution for non-integrated multiscale correlative light and electron microscopy. Journal of Structural Biology, 201(1), 63–75. https://doi.org/10.1016/j.jsb.2017.11.002

Liljeroos, L., Krzyzaniak, M. A., Helenius, A., & Butcher, S. J. (2013). Architecture of respiratory syncytial virus revealed by electron cryotomography. Proceedings of the National Academy of Sciences, 110(27), 11133–11138. https://doi.org/10.1073/pnas.1309070110

Liu, B., Xue, Y., Zhao, W., Chen, Y., Fan, C., Gu, L., Zhang, Y., Zhang, X., Sun, L., Huang, X., Ding, W., Sun, F., Ji, W., & Xu, T. (2015). Three-dimensional super-resolution protein localization correlated with vitrified cellular context. Scientific Reports, 5(1), 13017. https://doi.org/10.1038/srep13017

Lučić, V., Kossel, A. H., Yang, T., Bonhoeffer, T., Baumeister, W., & Sartori, A. (2007). Multiscale imaging of neurons grown in culture: From light microscopy to cryo-electron tomography. Journal of Structural Biology, 160(2), 146–156. https://doi.org/10.1016/j.jsb.2007.08.014

Lučić, V., Rigort, A., & Baumeister, W. (2013). Cryo-electron tomography: The challenge of doing structural biology in situ. The Journal of Cell Biology, 202(3), 407–419. https://doi.org/10.1083/jcb.201304193

Luque, D., & Castón, J. R. (2020). Cryo-electron microscopy for the study of virus assembly. Nature Chemical Biology, 16(3), 231–239. https://doi.org/10.1038/s41589-020-0477-1

Marko, M., Hsieh, C., Moberlychan, W., Mannella, C. A., & Frank, J. (2006). Focused ion beam milling of vitreous water: Prospects for an alternative to cryo-ultramicrotomy of frozen-hydrated biological samples. Journal of Microscopy, 222(1), 42–47. https://doi.org/10.1111/j.1365-2818.2006.01567.x

Marko, Michael, Hsieh, C., Schalek, R., Frank, J., & Mannella, C. (2007). Focused-ion-beam thinning of frozen-hydrated biological specimens for cryo-electron microscopy. Nature Methods, 4(3), 215–217. https://doi.org/10.1038/nmeth1014

Mastronarde, D. N. (2005). Automated electron microscope tomography using robust prediction of specimen movements. Journal of Structural Biology, 152(1), 36–51. https://doi.org/10.1016/j.jsb.2005.07.007

Moser, F., Pražák, V., Mordhorst, V., Andrade, D. M., Baker, L. A., Hagen, C., Grünewald, K., & Kaufmann, R. (2019). Cryo-SOFI enabling low-dose super-resolution correlative light and electron cryo-microscopy. Proceedings of the National Academy of Sciences, 116(11), 4804–4809. https://doi.org/10.1073/pnas.1810690116

Nahmani, M., Lanahan, C., DeRosier, D., & Turrigiano, G. G. (2017). High-numerical-aperture cryogenic light microscopy for increased precision of superresolution reconstructions. Proceedings of the National Academy of Sciences, 114(15), 3832–3836. https://doi.org/10.1073/pnas.1618206114

Oomens, A. G. P., Bevis, K. P., & Wertz, G. W. (2006). The Cytoplasmic Tail of the Human Respiratory Syncytial Virus F Protein Plays Critical Roles in Cellular Localization of the F Protein and Infectious Progeny Production. Journal of Virology, 80(21), 10465–10477. https://doi.org/10.1128/JVI.01439-06

Paul-Gilloteaux, P., Heiligenstein, X., Belle, M., Domart, M.-C., Larijani, B., Collinson, L., Raposo, G., & Salamero, J. (2017). eC-CLEM: Flexible multidimensional registration software for correlative microscopies. Nature Methods, 14(2), 102–103. https://doi.org/10.1038/nmeth.4170

Rigort, A., Bäuerlein, F. J. B., Leis, A., Gruska, M., Hoffmann, C., Laugks, T., Böhm, U., Eibauer, M., Gnaegi, H., Baumeister, W., & Plitzko, J. M. (2010). Micromachining tools and correlative approaches for cellular cryo-electron tomography. Journal of Structural Biology, 172(2), 169–179. https://doi.org/10.1016/j.jsb.2010.02.011

Sartori-Rupp, A., Cordero Cervantes, D., Pepe, A., Gousset, K., Delage, E., Corroyer-Dulmont, S., Schmitt, C., Krijnse-Locker, J., & Zurzolo, C. (2019). Correlative cryo-electron microscopy reveals the structure of TNTs in neuronal cells. Nature Communications, 10(1), 342. https://doi.org/10.1038/s41467-018-08178-7

Schellenberger, P., Kaufmann, R., Siebert, C. A., Hagen, C., Wodrich, H., & Grünewald, K. (2014). High-precision correlative fluorescence and electron cryo microscopy using two independent alignment markers. Ultramicroscopy, 143, 41–51. https://doi.org/10.1016/jultramic.2013.10.011

Schindelin, J., Arganda-Carreras, I., Frise, E., Kaynig, V., Longair, M., Pietzsch, T., Preibisch, S., Rueden, C., Saalfeld, S., Schmid, B., Tinevez, J.-Y., White, D. J., Hartenstein, V., Eliceiri, K., Tomancak, P., & Cardona, A. (2012). Fiji: An open-source platform for biological-image analysis. Nature Methods, 9(7), 676–682. https://doi.org/10.1038/nmeth.2019

Schorb, M., & Briggs, J. A. G. (2014). Correlated cryo-fluorescence and cryo-electron microscopy with high spatial precision and improved sensitivity. Ultramicroscopy, 143, 24–32. https://doi.org/10.1016/j.ultramic.2013.10.015

Schorb, M., Gaechter, L., Avinoam, O., Sieckmann, F., Clarke, M., Bebeacua, C., Bykov, Y. S., Sonnen, A. F.-P., Lihl, R., & Briggs, J. A. G. (2017). New hardware and workflows for semi-automated correlative cryo-fluorescence and cryo-electron microscopy/tomography. Journal of Structural Biology, 197(2), 83–93. https://doi.org/10.1016/j.jsb.2016.06.020

Schorb, M., Haberbosch, I., Hagen, W. J. H., Schwab, Y., & Mastronarde, D. N. (2019). Software tools for automated transmission electron microscopy. Nature Methods, 16(6), 471–477. https://doi.org/10.1038/s41592-019-0396-9

Schwartz, C. L., Sarbash, V. I., Ataullakhanov, F. I., Mcintosh, J. R., & Nicastro, D. (2007). Cryo-fluorescence microscopy facilitates correlations between light and cryo-electron microscopy and reduces the rate of photobleaching. Journal of Microscopy, 227(2), 98–109. https://doi.org/10.1111/j.1365-2818.2007.01794.x

Strauss, J. D., Hammonds, J. E., Yi, H., Ding, L., Spearman, P., & Wright, E. R. (2015). Three-Dimensional Structural Characterization of HIV-1 Tethered to Human Cells. Journal of Virology, 90(3), 1507–1521. https://doi.org/10.1128/JVI.01880-15

Strnad, M., Elsterová, J., Schrenková, J., Vancová, M., Rego, R. O. M., Grubhoffer, L., & Nebesářová, J. (2015). Correlative cryo-fluorescence and cryo-scanning electron microscopy as a straightforward tool to study host-pathogen interactions. Scientific Reports, 5(1), 18029. https://doi.org/10.1038/srep18029

Szeliski, R. (2011). Computer vision: Algorithms and applications. Springer.

Tang, G., Peng, L., Baldwin, P. R., Mann, D. S., Jiang, W., Rees, I., & Ludtke, S. J. (2007). EMAN2: An extensible image processing suite for electron microscopy. Journal of Structural Biology, 157(1), 38–46. https://doi.org/10.1016/j.jsb.2006.05.009

van Driel, L. F., Valentijn, J. A., Valentijn, K. M., Koning, R. I., & Koster, A. J. (2009). Tools for correlative cryo-fluorescence microscopy and cryo-electron tomography applied to whole mitochondria in human endothelial cells. European Journal of Cell Biology, 88(11), 669–684. https://doi.org/10.1016/j.ejcb.2009.07.002

Wan, W., Kolesnikova, L., Clarke, M., Koehler, A., Noda, T., Becker, S., & Briggs, J. A. G. (2017). Structure and assembly of the Ebola virus nucleocapsid. Nature, 551(7680), 394–397. https://doi.org/10.1038/nature24490

Wolff, G., Hagen, C., Grünewald, K., & Kaufmann, R. (2016). Towards correlative super-resolution fluorescence and electron cryo-microscopy: Towards super-resolution cryo-CLEM. Biology of the Cell, 108(9), 245–258. https://doi.org/10.1111/boc.201600008

Yi, H., Strauss, J. D., Ke, Z., Alonas, E., Dillard, R. S., Hampton, C. M., Lamb, K. M., Hammonds, J. E., Santangelo, P. J., Spearman, P. W., & Wright, E. R. (2015). Native Immunogold Labeling of Cell Surface Proteins and Viral Glycoproteins for Cryo-Electron Microscopy and Cryo-Electron Tomography Applications. Journal of Histochemistry & Cytochemistry, 63(10), 780–792. https://doi.org/10.1369/0022155415593323

Zachs, T., Schertel, A., Medeiros, J., Weiss, G. L., Hugener, J., Matos, J., & Pilhofer, M. (2020). Fully automated, sequential focused ion beam milling for cryo-electron tomography. ELife, 9, e52286. https://doi.org/10.7554/eLife.52286

Zhang, P. (2013). Correlative cryo-electron tomography and optical microscopy of cells. Current Opinion in Structural Biology, 23(5), 763–770. https://doi.org/10.1016/j.sbi.2013.07.017

