## Supplementary figures for "CorRelator: An interactive and flexible toolkit for high-precision cryo-correlative light and electron microscopy"

### Supplementary Figure 1

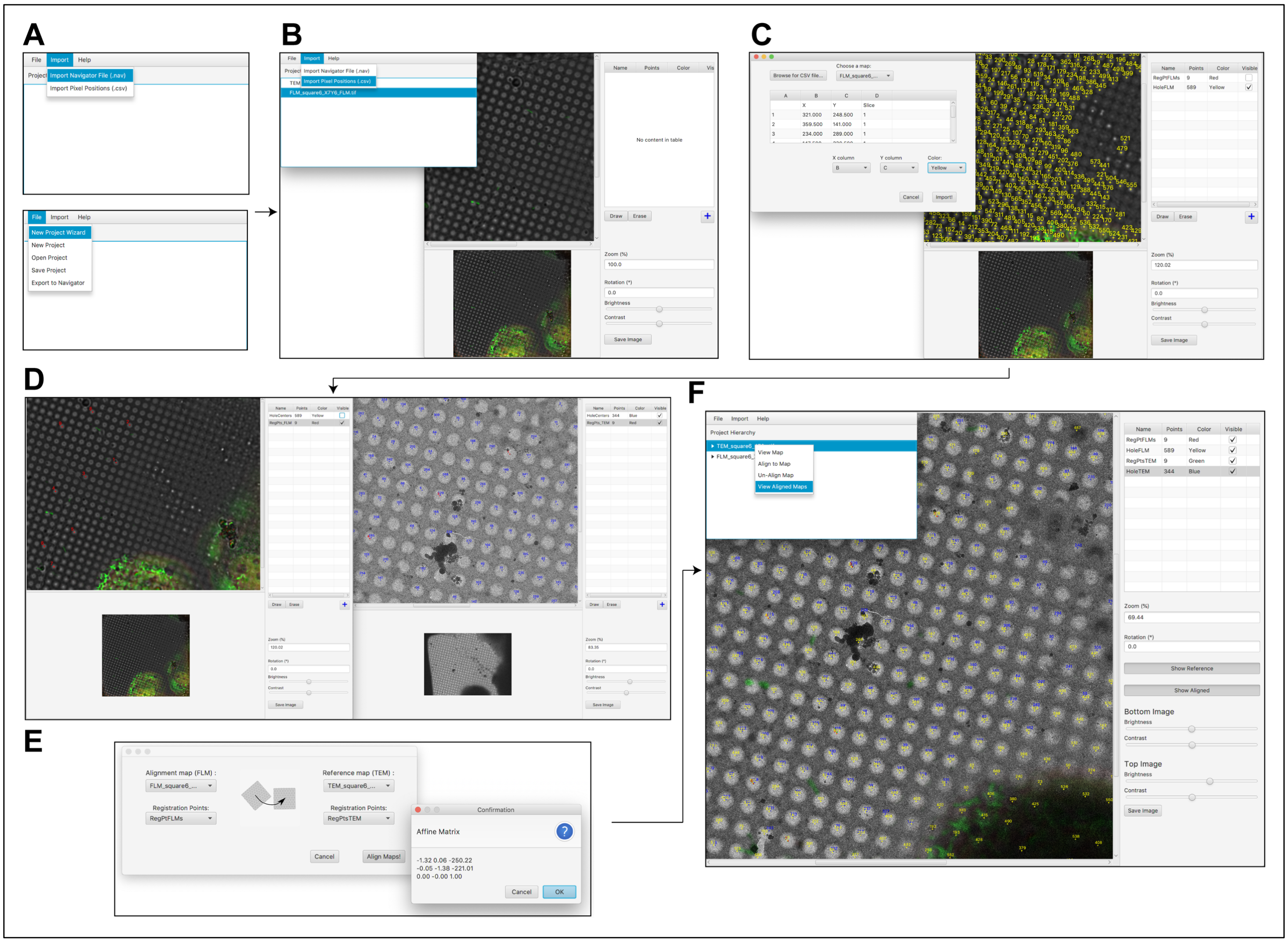


**Supplementary Fig. 1.** Six steps in CorRelator guide the user through interactive on-the-fly registration and correlation. (A) The user starts by loading a baseline SerialEM navigator file Nav_1 through the *Wizard* or main *Project* View. (B) In step 2, the user can choose to import independent coordinate csv files associated with certain FLM and TEM map items in Nav_1 and interactively edit (add or delete) points in individual image viewer windows. Each csv file is listed as a group of points. (C) Third, the matching reference point pairs for registration can be specified using either existing imported external points for guidance or newly assigned markers. (D-E) After the registration points are chosen, the transform matrix M is calculated. (F) The alignment performance can be quickly assessed through the overlay of correlated cryo-FLM and EM images based on the calculated M. The user can un-align two images and re-initiate the registration and transformation process. All operations and calculated matrices can be saved as a project that can be continued later. A new functional navigator file Nav_2 can be written out and re-loaded back in SerialEM.

**Supplementary Figure 2**

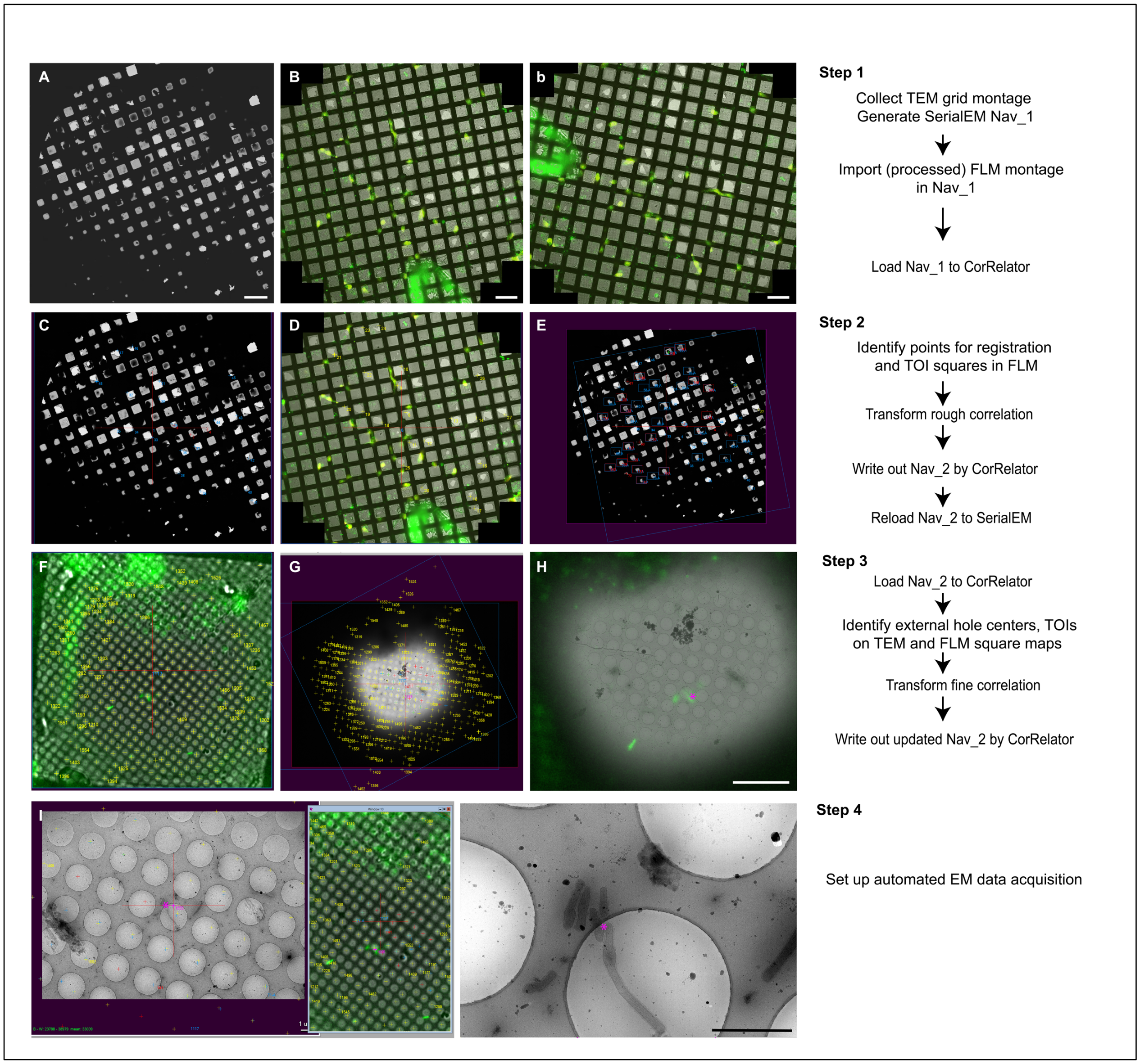


**Supplementary Fig.2** Four steps of CorRelator cryo-CLEM workflow demonstrated by representative images and SerialEM screenshots. The procedure to set up the rough correlation on the whole grid level (A-E, Step 1-2) and subsequent fine correlation using reference hole centroids and RSV particles as targets of interest (TOI) (F-J, Step 3-4). (A) Full cryo-EM grid map is acquired by SerialEM. (B) Cryo-FLM mosaic of the same whole grid B is edited by flipping horizontally and rotating the raw mosaic (b) clockwise ~80 º, based on (A), prior to its import into

SerialEM. Fluorescent signal from RSV-infected cells (red) and the RSV F glycoprotein on RSV-infected cells or released particles (green). (C-D) Screenshots of the TEM and FLM maps reloaded from the Nav-2 file. Externally picked landmark points (blue in the TEM map and yellow in the FLM map) have updated stage positions. (E) Screenshot of post-correlated TEM grid map. Blue and red boxes indicate the positions where targeted squares are recorded based on rough correlation. (F-G) Screenshots of square FLM image and corresponding correlated TEM maps where viral particles are green. The hole center stage positions display as yellow dots in the FLM map (F) and as yellow and blue dots in the post-transformed correlated TEM map (G). (H) Superposition of the TEM and FLM images after correlation. The TOI was identified by the pink asterisk (numbered as the point item 1555 in SerialEM). (I) Screenshot of the magnified fine correlation. (J) Magnified view of the cryo-EM image based on the transformed position of the matching TOIs from (H and I). Image rotation between H and I resulted from magnification lens change under TEM. Scale bars: 200 µm in A, B, b, 10 µm in H, and 1 µm in J.

**Supplementary Figure 3**

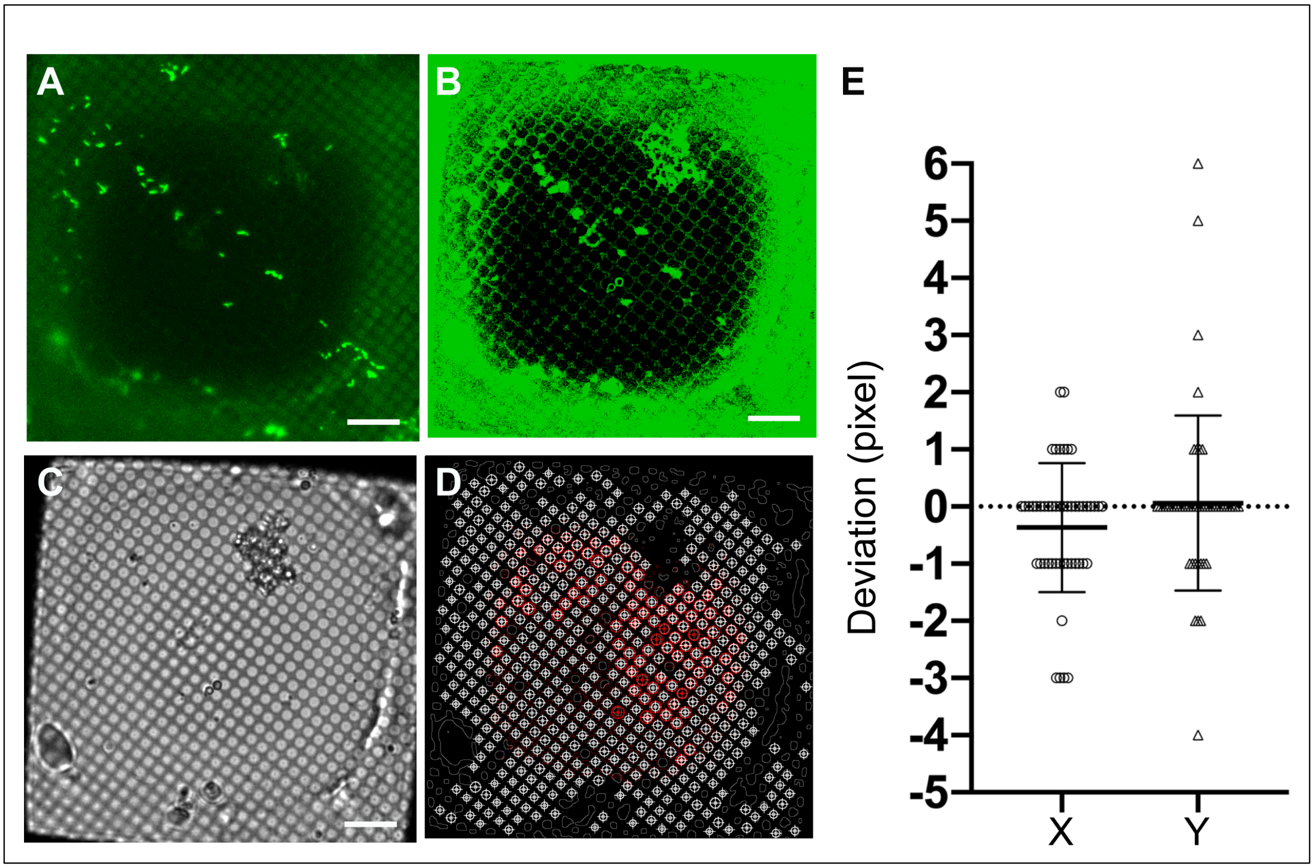


**Supplementary Fig. 3** Comparison of hole centroid identification in the fluorescent and bright field frames. (A) Vitrified immuno-labeled RSV viral particles (green, GFP) is imaged with Leica EM cryo-CLEM system in the GFP (emission 525 nm) channel at 50x (130 nm/pixel). (B) Enhanced version of A shows the detectability of the holes in the carbon film after filtering. (C) Cryo-CLEM bright field image of the same recorded area goes through the same algorithmic method for hole identification. (D) Overlay of automated identified hole centroid coordinates using enhanced fluorescent image (B, red cross coordinates) and bright field image (C, white cross coordinates) shows the deviation between two channels. (E) The deviation in pixels of the hole center position identified in the bright field image and in the enhanced fluorescent image ($n = 50$) is quantified. Scale bars = 10 $\mu$m.

**Supplementary Figure 4**

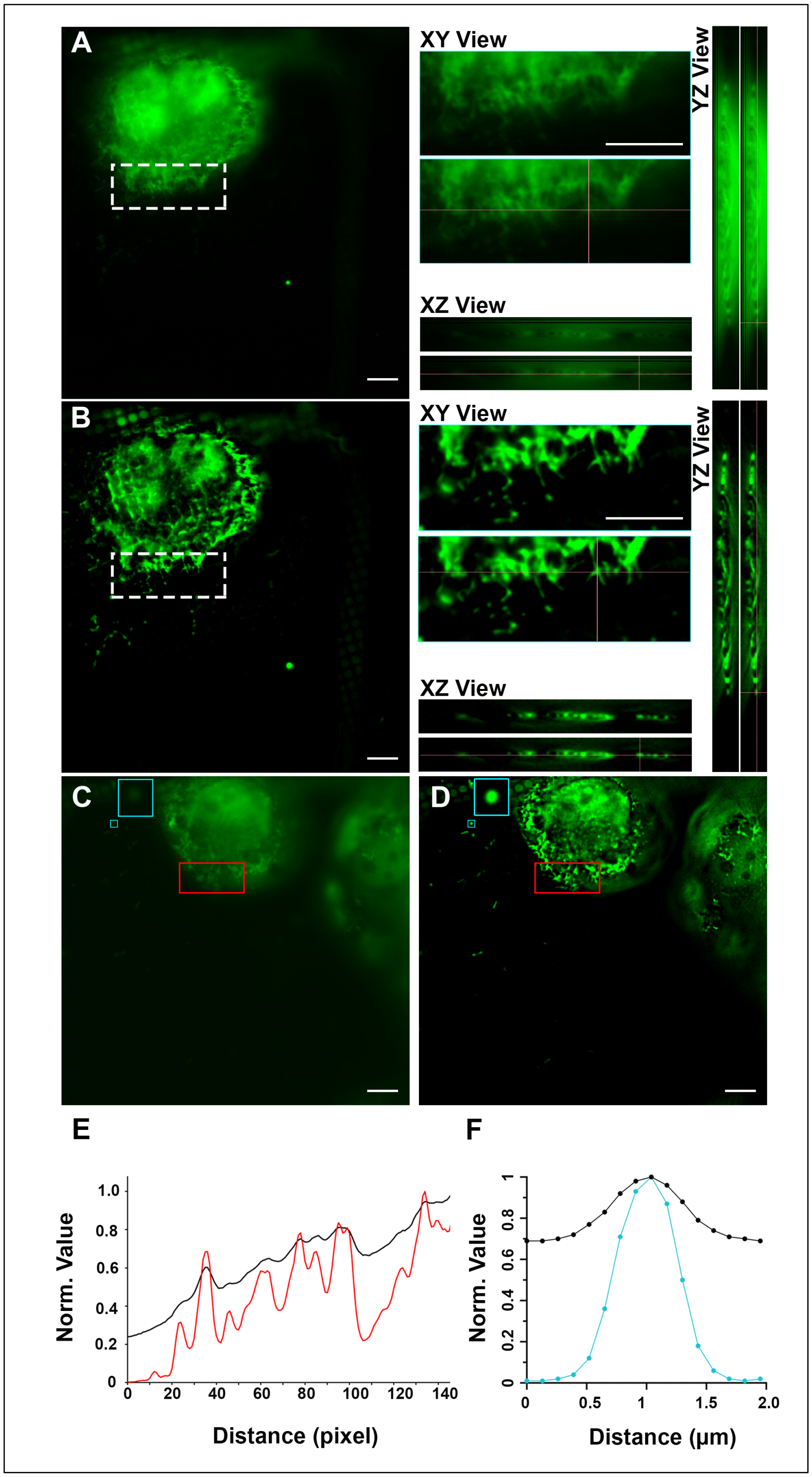

**Supplementary Fig. 4.** Improved contrast and image resolution after Small Volume Computational Clearance (SVCC). (A) A single plane unprocessed fluorescent image is extracted from a z-stack imaged on the Leica EM cryo-CLEM system, in green channel. XY, XZ, and YZ orthogonal views of the selected region in the dashed white box are shown. (B) The resultant image after SVCC of the raw data (A) displays sharp and clean features in XY, XZ, and YZ views. (C, D) A second representative image of pre- (C) and post-SVCC (D) are compared. (E) Normalized X-axis intensity plot file of the red boxed area in pre-SVCC frame C (black line) and post-SVCC frame D (red line) show matching intensity peaks, suggesting undetectable pixel changes after SVCC. (F) Representative normalized X-axis intensity measurement of a blue-boxed (zoomed view on the right side) single 500 nm TetraSpeck-bead in C (black line) and D (red line), contributes to measurement of the lateral PSF of the pre- and post-SVCC images. Scale bars = 10 µm.

**Supplementary Figure 5**

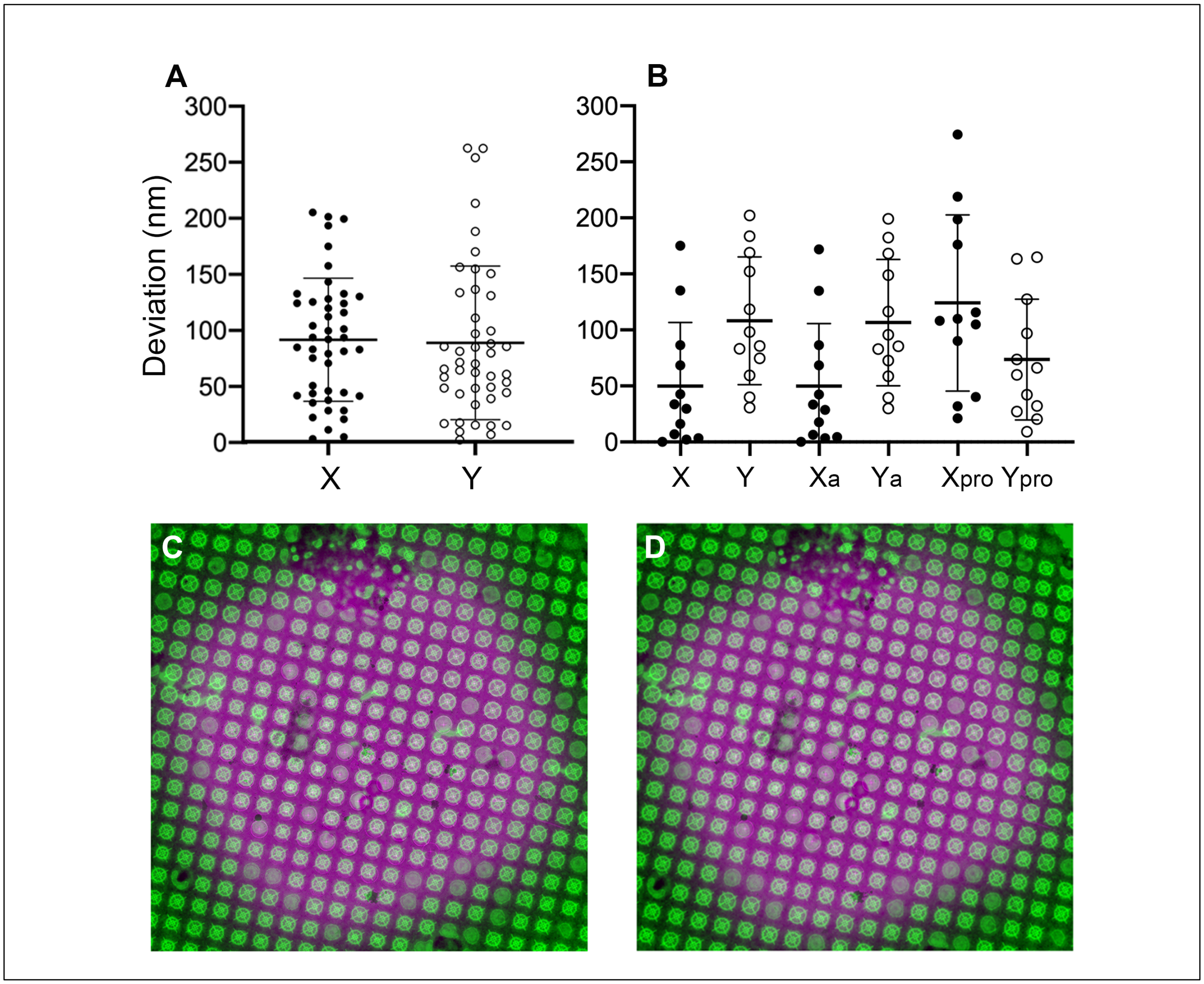


**Supplementary Fig. 5**. Evaluation of accuracy localization error in post-acquisition correlation. (A) Relocation errors in X and Y axis by CorRelator (n = 44). The mean is 68.6 nm and 66.8 nm in X and Y with the deviation of 41.3 nm and 52.2 nm in X and Y, respectively. (B) Predicted errors in X and Y by CorRelator, MATLAB affine and MATLAB projective transformation (n = 12). The comparison was done using the same pairs of registration points and the same fluorescent targets of interest. X and Y for CorRelator (standard deviation of 56.6 nm and 57.0 nm, respectively), Xa and Ya for MATLAB affine transformation (standard deviation of 55.7 nm and

56.4 nm, respectively), Xpro and Ypro for MATLAB projective transformation standard deviation of (56.4 nm and 78.7 nm, respectively). (C) Superimposition of the TEM image (green) and transformed fluorescent image (purple) using MATLAB affine transformation. (D) Superimposition of the TEM image (green) and transformed fluorescent image (purple) using MATLAB projective transformation.

**Supplementary Figure 6**

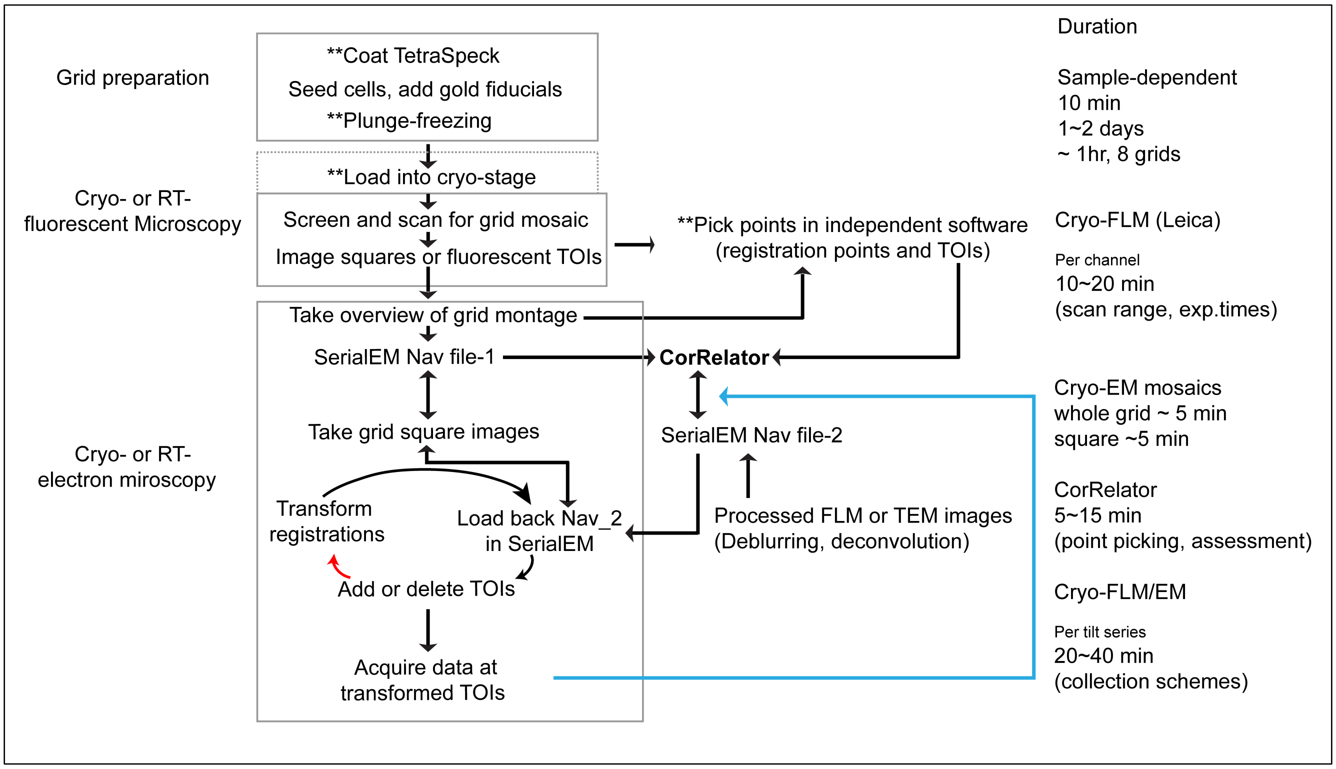


**Supplementary Fig. 6.** CorRelator enables flexible and smooth correlative light and electron microscopy (CLEM) workflows for ambient and cryogenic operations. The black and red arrow flows illustrate two flexible on-the-fly correlation approaches supported by CorRelator corresponding to Route ① and ② in Figure 1, respectively. The blue arrow indicates CorRelator-mediated post-acquisition TEM-FLM correlative mapping with collected data. The grey boxes note critical sections of the correlation experiment. Double arrows show that the workflow can go both ways. Both SerialEM navigator file-1 and -2 (Nav_1 and Nav_2) can be updated at the end of the individual steps. For example, after Nav_2 guides the TEM square map collection at the stage of rough correlation, it will be re-used by CorRelator for fine correlation, thus becoming the new Nav_1. Double asterisks (**) signify optional steps, e.g., TetraSpeck addition, cryogenic operations including plunge-freezing and loading into the cryo-stage, or the import of external coordinates through csv files from independent software. The estimated time to complete each step is listed on the right.

**Supplementary Figure 7**

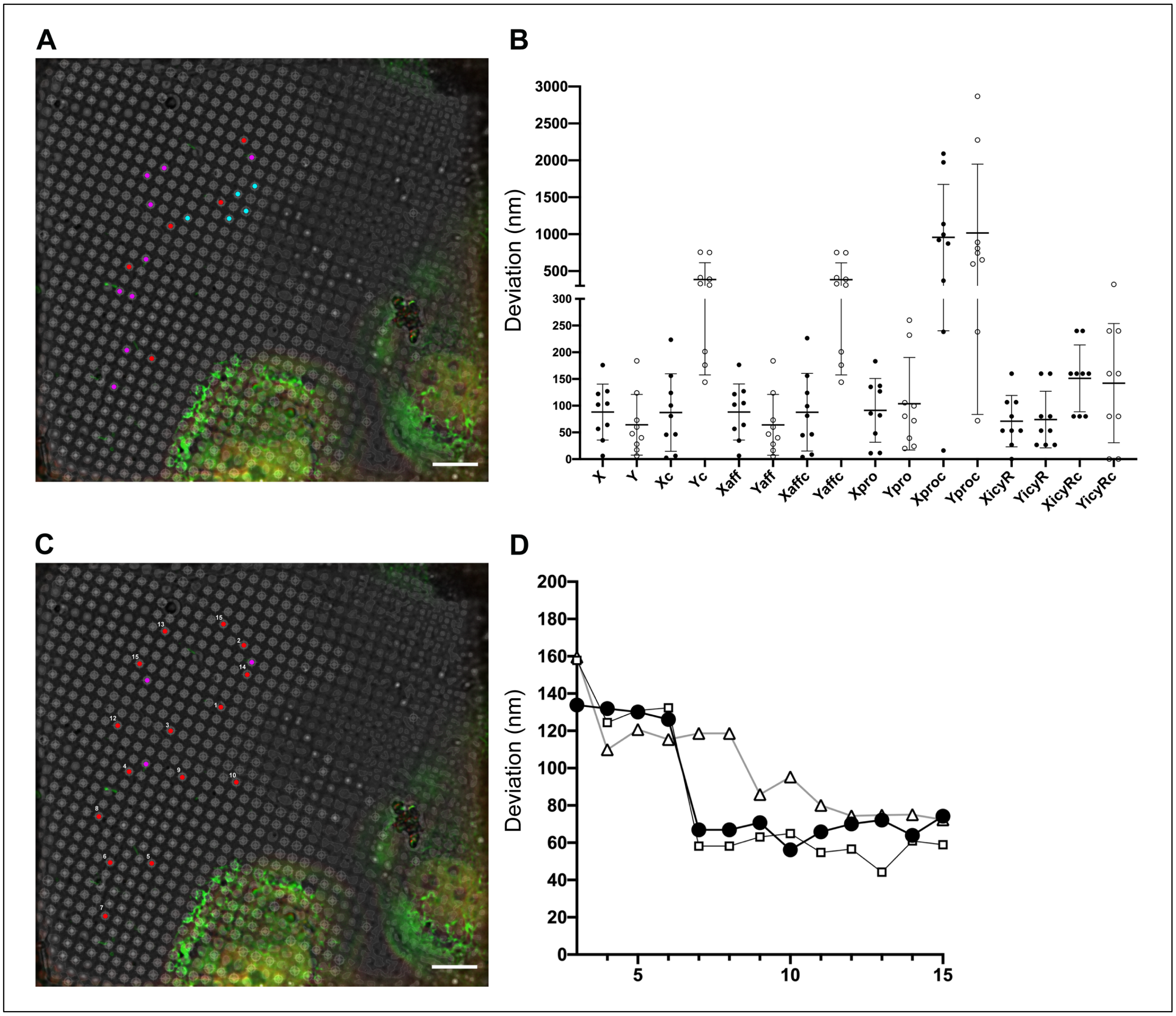


**Supplementary Fig. 7.** Comparison and prediction error monitor. (A) Five registration points and nine target points (leave-one-out method) picked on a 2D cryo-FLM image of vitrified cells and used for comparison of predicted errors shown in (B). Spreading reference points are red (n = 5); locally centered reference points are cyan (n = 5); target points are magenta (n = 9). (B) Comparison of predicted errors of the picked targets (purple, A) in X and Y by several software tools. From left to right: X/Y and Xc/Yc are errors predicted by CorRelator using spreading and locally centered registration points, Xaff/Yaff and Xaffc/Yaff by MATLAB affine transformation, Xpro/Ypro and Xproc/Yproc by MATLAB projective transformation, XicyR/YicyR and XicyRc/YicyRc by eC-CLEM (ICY package) transformation. (C) Spreading reference points (red, numbered) and targets (magenta) picked to monitor the predicted error as the reference points increase. (D) Predicted errors on monitored three magenta targets (A). The X axis is number of points used for the reference, labeled (C). Scale bars = 10 μm.

**Supplementary Movie 1.** This movie demonstrates the use of CorRelator for correlative cryo-FLM and automated cryo-ET data collection. Thirteen steps are presented as screenshots from active GUI windows in CorRelator and SerialEM. The last frame is the complete set of cryo-FLM targets used for automated cryo-ET data collection.

**Supplementary Table 1.** Comparison of available software or scripting options for image registration and correlation. References are initial software development and use.

| Software | Availability | Application | Requirement | Output Ready for TEM | Registration | Error Prediction | User Imported points | References |
|---|---|---|---|---|---|---|---|---|
| **CorRelator** | Free, source code available | 2D-2D On-the-fly, Post-acquisition cryo-CLEM | Java | Yes | Manual & assisted manual | Yes | Yes | N/A |
| **MATLAB Plugin Package** | Licensed under MATLAB | 2D-2D, Post-acquisition cryo- & plastic CLEM | MATLAB | No | Manual | Yes | No | (Kukulski et al. 2011); |
| **MATLAB Script** | Licensed under MATLAB | 2D-2D, On-the-fly, cryo-CLEM | MATLAB | Yes | Manual | Yes | No | (Fu et al. 2019) |
| **eC-CLEM** | Free, source code available | 2D-2D, 3D-3D, Post-acquisition cryo- & plastic CLEM | Java, Icy | No | Manual & Automatic | Yes | No | (Paul-Gilloteaux et al. 2017) |
| **PIE-scope** | Free, source code available | 2D-2D, on-the-fly cryo-FLM-FIB/SEM | Python, Autoscript (FEI microscope systems, licensed) | No | Manual | No | No | (Gorelick et al. 2019) |
| **ColorView** | Licensed under LabVIEW | 2D-2D, on-the-fly cryo-CLEM | LabVIEW | N/A | Manual | No | No | (Li et al. 2018) |
| **SerialEM** | Free | 2D-2D, on-the-fly cryo-CLEM | Windows | Yes | Manual | No | Yes | (Schwartz et al., 2007) |
| **TurboReg** | Free, source code available | 2D-2D | Java, ImageJ | No | Manual | No | No | (Keene et al. 2014) |

**Appendix-A**

The Navigator module is one of the key functions in SerialEM to save entry metadata, link microscope stage positions and image coordinates, mark positions of interest on map images, guide the stage movements, and to track and reload map images through relative paths. The Navigator maintains a list of item entries including points and maps. A map is a specialized single-frame image or multi-tile montage that serves as a canvas and defines an area with its four corner stage positions. A point is a single stage location made on a map. Users start a Navigator file by generating a map that could be collected at a suitable magnification. Targets of interest are made as single points on one map. Under each item are an organized block of sections, among which values of MapScaleMat, RawStageXYZ, and MapHeightWidth store pixel-to-stage-position matrix information to transform points in pixel positions made on an image to corresponding stage positions. The transformation $M_{stage2pixel}$ is determined by stage shift calibration (the relationship between stage movements and position on the camera at a particular magnification) in SerialEM, and reported in the map item entries of PtsX and PtsY (X- and Y- stage coordinates of points), MapScaleMat (Stage to pixel scale and rotation matrix for drawing, based on pixels of initial map image), MapWidthHeight (size of initial map image in pixels) and RawStageXY (raw stage position before adjustment for translation). The 2D transformation must fit in:

$$\begin{bmatrix} P_{ts}X - RawStageX \\ P_{ts}Y - RawStageY \end{bmatrix} * MapScaleMat + \begin{bmatrix} 0.5MapWidth \\ 0.5MapHeight \end{bmatrix} = \begin{bmatrix} P_x \\ P_y \end{bmatrix}$$

Where $\begin{bmatrix} P_{ts}X \\ P_{ts}Y \end{bmatrix}$ is the stage position and $\begin{bmatrix} P_x \\ P_y \end{bmatrix}$ is the corresponding image coordinate pair.
